# Localized Delivery of Growth Factors from Microparticles Modulate Osteogenic and Chondrogenic Gene Expression in Growth Factor-dependent Manner in an *ex vivo* Chick Embryonic Bone Model

**DOI:** 10.1101/2024.12.05.627026

**Authors:** Hassan Rashidi, Helen C. Cox, Omar Qutachi, Dale Moulding, Lisa J. White, Emma L. Smith, Janos M. Kanczler, Luis Rojo, Michael Rotherham, James R. Henstock, Molly M. Stevens, Alicia J. El Haj, Richard O.C. Oreffo, Kevin M. Shakesheff, Felicity R.A.J. Rose

## Abstract

Growth factors play a crucial role in regulating various cellular functions, including proliferation and differentiation. Consequently, the biomaterial-based delivery of exogenous growth factors presents a promising strategy in regenerative medicine to manage the healing process and restore tissue function. For effective therapeutic applications, it is essential that these active compounds are precisely targeted to the site of regeneration, with release kinetics that align with the slow pace of tissue growth. We have developed an ex vivo model utilizing a developing embryonic chick bone, and using PLGA based microparticles as controlled-release systems, allowing for the investigation of spatiotemporal effects of growth factor delivery on cell differentiation and tissue formation. Our findings demonstrate that BMP2 and FGF2 can significantly alter cell morphology and zonally pattern collagen deposition within the model, but only when the growth factor presentation rate is carefully regulated. Furthermore, the growth factor-dependent responses observed underscore the potential of this model to explore the interactions between cells and the growth factors released from biomaterials in an approach which can be applied for bone tissue engineering.

## 1. Introduction

Bone regenerative therapeutics focus on the restoration of osseous defects and in particular critical size defects resulting from revision surgery, osteonecrosis, pathological deformation, traumatic injuries, infection and tumor resection (Shrivats et al., 2014). While bone grafting is regarded as the gold standard procedure of choice, limited donor site availability and donor site morbidity remain significant challenges of autologous grafting. Furthermore, rejection and infection are considered as major drawbacks of allogenic bone grafts (Dimitriou et al., 2011). This has heightened the demand for synthetic tissue-engineered alternatives to address complex clinical conditions, such as non-union fractures. Scaffolds with desired mechanical and physical properties have been considered as alternative approaches to facilitate the regeneration process. However, lack of functional moiety to promote interaction with human progenitor cells and insufficient biomimetic properties are major challenges that must be addressed to promote functional tissue formation using simple polymeric biomaterials (Khan and Tanaka, 2017). To overcome these limitations, strategies that integrate cell and growth factor delivery with biomaterial scaffolds have shown promise in achieving superior regenerative outcomes (Mehta et al., 2012, Wei et al., 2022).

To date, various growth factors have been explored for their potential applications in bone and cartilage tissue engineering, including bone morphogenic protein 2 (BMP2) (Johnson et al., 1988, Welch et al., 1998, Kandziora et al., 2002), bone morphogenic protein 7 (BMP7) (Ripamonti et al., 1996), transforming growth factor 1 (TGFβ1) (McKinney and Hollinger, 1996, Beck et al., 1998), transforming growth factor 2 (TGFβ2) (Critchlow et al., 1995), transforming growth factor 3 (TGFβ3) (Steffen et al., 2001), platelet-derived growth factor (PDGF-BB) (Nash et al., 1994), vascular endothelial growth factor (VEGF) (Kaigler et al., 2006), and fibroblast growth factor 2 (FGF2) (Radomsky et al., 1999) have been examined for bone tissue engineering. A substantial portion of research in this field has concentrated on potential therapeutic application of BMP2 (Gothard et al., 2014, Zhu et al., 2022). While significant improvement in healing has been reported following BMP2 delivery at bone defect or fracture sites, several adverse effects have been reported, including ectopic bone formation, osteolysis, retrograde ejaculation and greater risk of malignancy (Deutsch, 2010, Carragee et al., 2011, Liu et al., 2023). It has been speculated that these adverse effects may be linked to the delivery of supraphysiological concentrations of BMP2 due to insufficient or absent sustained release mechanisms (Deutsch, 2010).

To enhance efficacy while reducing dosages, the field has shifted toward the manufacturing of scaffolds incorporating a lower quantity of bioactive cargo with sustained release over time (Simmons et al., 2004, Tayalia and Mooney, 2009). To achieve the desired therapeutic outcomes, various sophisticated carriers have been manufactured for bone tissue engineering to enable spatiotemporal delivery of single or indeed multiple growth factors and recombinant morphogens (Mehta et al., 2012). The effect of a growth factor is governed by various factors including; i) concentration of the growth factor, ii) the cell type and the receptors involved, and iii) the intracellular transduction pathway at play (Lee et al., 2011). Despite recent advances in fabrication of growth factor-loaded materials, lack of appropriate biological *in vitro* assays to assess their release profile and efficacy has remained as one of the main obstacles (Bock et al., 2016). A number of approaches have been developed to test manufactured polymeric systems; including, *in vitro* osteoinduction assays (Basmanav et al., 2008, Yilgor et al., 2010, Fois et al., 2024) and subcutaneously implanted diffusion chambers in SCID mice (Huang et al., 2005). However, these approaches lack the complexity to fully delineate the therapeutic potential of developed growth factor-loaded polymers in bone tissue engineering. Therefore, a more physiologically-relevant model is highly desirable to accurately assess the efficacy and usefulness of polymeric carriers containing single, dual or multiple growth factors.

Bone development in embryonic chicks follows a similar developmental pathway as human bone during endochondral ossification (Kanczler et al., 2012, Smith et al., 2012). making embryonic chick bones a useful model for bone regeneration. In this study, the organotypic chick embryonic bone was developed as a more physiologically-relevant model to analyze the localized effect of BMP2 released from PLGA-based MPs (BMP2-MPs) with 10% w/w PLGA-PEG-PLGA tri-block copolymer as plasticizer and average diameter of 20 μm alongside appropriate controls. Additionally, basic fibroblast growth factor (FGF2)-loaded MPs (FGF2-MPs) were injected to evaluate cellular response to a non-osteoinductive growth factor, with the aim of determining growth factor-dependency of cellular response in the chick embryonic hyaline cartilage.

## 2. Materials and methods

### 2.1. Microparticle manufacturing and growth factor encapsulation

Poly(vinyl alcohol) (PVA, molecular weight: 13000 – 23000 Da, 87-89% hydrolyzed), human serum albumin (HSA), Poly (DL-lactide-co-glycolide) (PLGA) polymers with lactide: glycolide ratios of 50:50 (DLG 4.5A 59 kDa) were purchased from Evonik Industries (Alabama, USA). Recombinant human BMP2 (BMP2) was obtained from 2 different sources including Professor Walter Sebald (University of Wurzburg, Germany) and ORF Genetics (BMP2^ORF^, ORF Genetics Ltd., Iceland). Recombinant human FGF2 was purchased from ORF Genetics (ORF Genetics Ltd., Iceland). PLGA-PEG-PLGA tri-block was produced in house following protocol published previously (Kirby et al., 2011). PLGA-based MPs were manufactured using a water-in-oil-in-water (w/o/w) double emulsion method and characterized as previously described (Kirby et al., 2011). Briefly, PLGA-PEG-PLGA tri-block copolymer was added to PLGA to provide weight percentages of 10% (w/w) of the 1g total mass in 5 ml dichloromethane. BMP-2 and 1% human serum albumin (HSA, Sigma-Aldrich, UK) solution were prepared at a ratio of 1:9 for a 1% w/w loading in the MPs. In order to manufacture MPs, the aqueous solution of 1% HSA and BMP-2 was added to a solution of PLGA-tri block copolymer. These phases were homogenized for two minutes at 4,000 rpm in a Silverson L5M homogenizer (Silverson Machines, UK) to form the water-in-oil emulsion. This primary emulsion was transferred to 200 ml 0.3% (w/v) polyvinyl alcohol (PVA) solution and was homogenized for a second time at 9,000 rpm. The resultant double emulsion was stirred at 300 rpm on a Variomag 15-way magnetic stirrer for a minimum of 4 hours to facilitate DCM evaporation, then washed and lyophilized (Edwards Modulyo, IMA Edwards, UK). The MPs were stored at -20 °C until injection. FGF2-MPs were manufactured following the same procedure. Fluorescently-labelled MPs were manufactured by addition of 0.001% Coumatin-6 (Sigma-Aldrich, UK) as described previously (Qutachi et al., 2013).

### 2.2. Microparticle characterization

Microparticles were characterized as described elsewhere (Kirby et al., 2011). Briefly, a suspension of the microparticles was prepared in double deionized water and sized using a laser diffraction method to measure the size distribution of microparticles (Supplementary Figure 1, Coulter LS230, fitted with the hazardous fluids module, Beckman Coulter, UK) while under agitation to prevent the particles settling. In order to measure the encapsulation efficiency of protein within microparticles, 10 mg microparticles were incubated in 750 µl dimethylsulphoxide (DMSO) at room temperature for 1 hour followed by the addition of 150 µl of 0.5% SLS/0.02 N NaOH for a further incubation at room temperature for another hour. Protein concentration of the resulting solution was measured using a bicinchoninic acid (BCA) protein assay kit (Thermo Scientific) and compared against a standard curve of HSA conducted at the same time. 150 µl of sample were mixed with 150 µl of BCA and incubated for 2 hours at 37 °C. The absorbance at 462 nm was measured using a Tecan infinite 200 plate reader. To study the release profile, aliquots of the microparticles (100 mg) were suspended in 3 ml of phosphate-buffered saline (PBS) and then incubated on an orbital shaker set at 5 rpm at 37 °C. The PBS was completely replaced at regular intervals and the protein content was measured using the BCA protein assay kit (Supplementary Figure 2). Calibration standards reflected the formulation of protein within the microparticles and were prepared in PBS.

### 2.3. Microinjection

GF-loaded MPs were resuspended in PBS and were injected into various sites in the femur (both cartilaginous epiphyses and the mid-point of the diaphyseal bone collar) using a Femtojet system (Eppendorf, Germany) and glass capillary needles with a ∼50-400 μm tip diameter (Supplementary Figure 3) pulled using a Flaming/Brown Micropipette Puller (Sutter, USA). Roughly 20 nl of material was injected into each site under sterile conditions with the aid of a dissecting microscope. Following optimization of the injection procedure, FGF2-MPs and BMP2-MPs were injected into chick embryonic bones. FGF2 solution, BMP2 solution and other appropriate controls including 1% HSA solution (carrier protein), and blank MPs (containing only HSA) were also injected following a similar procedure. In order to facilitate monitoring of the injection procedure under the microscope, GF-loaded MPs were mixed with labelled blank MPs in a ratio of 5:1. Endotoxin free BMP2-loaded MPs (BMP2^ORF^-MPs) were injected following a similar procedure to rule out the potential effect of endotoxin.

### 2.4. Organotypic culture of embryonic chick bones

Following injection, organotypic cultures of embryonic day 11 (E11) chick bones were performed as described previously (Kanczler et al., 2012). Briefly, injected bones were placed onto Millicell inserts (0.4-µm pore size, 30-mm diameter; Millipore) in Falcon^TM^ six-well tissue culture plate (Becton Dickinson Labware, USA) containing 1 ml minimum essential medium alpha (α-MEM) (Gibco) containing 100 units penicillin, 100 µg/ mL streptomycin, and 100 µM L-ascorbic acid 2-phosphate (Sigma-Aldrich) per well at the liquid/gas interface. Bones were cultured for 10 days at 37 °C, 5% CO_2_ with media changed every 24h.

### 2.5. Histological Examination

Bones were fixed in 4% paraformaldehyde for at least 24h. Samples were processed through a series of graded ethanol, cleared in xylene and embedded in low-melting point paraffin wax. Ten micrometer sections were cut and stained with Alcian blue/Sirius red staining to assess proteoglycan and collagen contents as described previously (Kanczler et al., 2012). Briefly, sections were first deparaffinized and rehydrated through graded alcohols and water. Then, sections were stained with Weigert’s Hematoxylin and differentiated in acid/alcohol followed by staining with 0.5% Alcian blue for proteoglycan-rich cartilage matrix and 1% Sirius red for collagenous bone matrix. Subsequently, sections were dehydrated and cleared before mounting in DPX. Images were captured using the Nanozoomer Digital Pathology (Hamamatsu Photonics K.K., Japan).

### 2.6. Immunohistochemical (IHC) Staining

Bone and cartilage markers were assessed by analyzing expression of collagen type I (COL-I), type II (COL-II), type X (COL-X) and osteocalcin (OC) as described previously (Smith et al., 2012). Briefly, deparaffinized and rehydrated sections were treated in tri-sodium citrate buffer (Fisher, UK) for antigen retrieval. Then samples were washed in running water and incubated with 3% hydrogen peroxide to quench endogenous peroxidize activity. Sections were digested with hyaluronidase for COL-II immunostaining. Non-specific binding was blocked with 1% bovine serum albumin (BSA) followed by overnight incubation at 4 °C with primary antibodies diluted appropriately in 1% BSA. Sections were subsequently washed in PBST (PBS + 0.5% Tween 20) and incubated with biotinylated horse anti-rabbit IgG secondary antibody for COL-I and COL-II and biotinylated horse anti-mouse IgG secondary antibody (Vector Laboratories Inc) for OC diluted in 1% BSA. ExtrAvidin^®^ Peroxidase and 3-amino-9-ethylcarbazole (AEC) substrate solution (Sigma Aldrich) was used to visualize brown immune complex reaction product, with Alcian blue counter stain. Negative controls, either exclusion of only primary antibody or both primary and secondary antibodies, showed absence of any positive staining.

### 2.7. Environmental Scanning Electron Microscopy

Ten micrometer sections were cut and mounted on Thermanox^TM^ plastic coverslips (Nunc, USA). Then sections were deparaffinized and rehydrated through graded alcohols and water. Images were obtained using an environmental scanning electron microscope (Philips FEG ESEM) equipped with a Trecor detector set at an accelerating voltage of 10 kV under a vacuum of 4.0-5.8 Torr and 6-10% humidity.

### 2.8. Image Analysis

Images were captured using the Nanozoomer Digital Pathology (Hamamatsu Photonics K.K., Japan). The intensity was assessed by three-level thresholding of the COL-I staining of 6 independent samples using Image J software and a purpose-built macro (https://github.com/DaleMoulding/Fiji-Macros?tab=readme-ov-file#3-level-dab-quantifier).

Image analysis was performed on sections obtained from independent microinjections and sections from the same injections. In case of Blank-MP injected samples, image analysis was performed on four independent injections and at least two sections from each injection (total n=21). Analysis on BMP2-MPs injected to PZ was performed on four independent injections and two sections of one injection (PZ1 and 2) and one sections of three other injections (PZ 3, 4 and 5; total n=5). For BMP2-MPs injected to CZ, analysis was performed on five independent injections and two sections of three samples (CZ1 and 2, CZ4 and 5 and CZ7 and 8) and one sections of two other injections (CZ 3 and CZ6; total n=8).

### 2.9. Statistical Analysis

Data were analyzed by GraphPad Prism (version 10). We did not use statistical methods to predetermine sample size, there was no randomization designed in the experiments, and the studies were not blinded. Significance was determined by One-way analysis of ANOVA with Tukey’s multiple comparisons test with a single variance. Data are represented as mean ± SEM, \**p* < 0.05, \*\**p* < 0.01, \*\*\**p* < 0.001. \*\*\*\**p* < 0.0001.

## 3. Results

### 3.1. Injection procedure

To achieve a minimally invasive injection, a range of glass needles with external diameters of 50-400 μm were initially tested (Supplementary Figure 3). Following injection, embryonic chick bones injected with fluorescently-labelled MPs were sectioned and imaged. As minimal-invasive injection and retention of injected MPs in close proximity to host cells was of utmost importance to accurately evaluate the effects of released GFs, samples were examined microscopically for evidence of damage to tissue architecture surrounding injection sites and cell localization around the injected MPs. The results indicated that minimal damage with negligible alteration to the architecture of native tissue could be achieved using needles with defined narrow tips and an external diameter of ∼100 μm. Although the volume and the number of injected MPs were increased 3-4 fold with the use of glass needles with wider bores, injection with such needles resulted in substantial tissue damage and the formation of large cavities (Figure 1).

**Figure 1:**
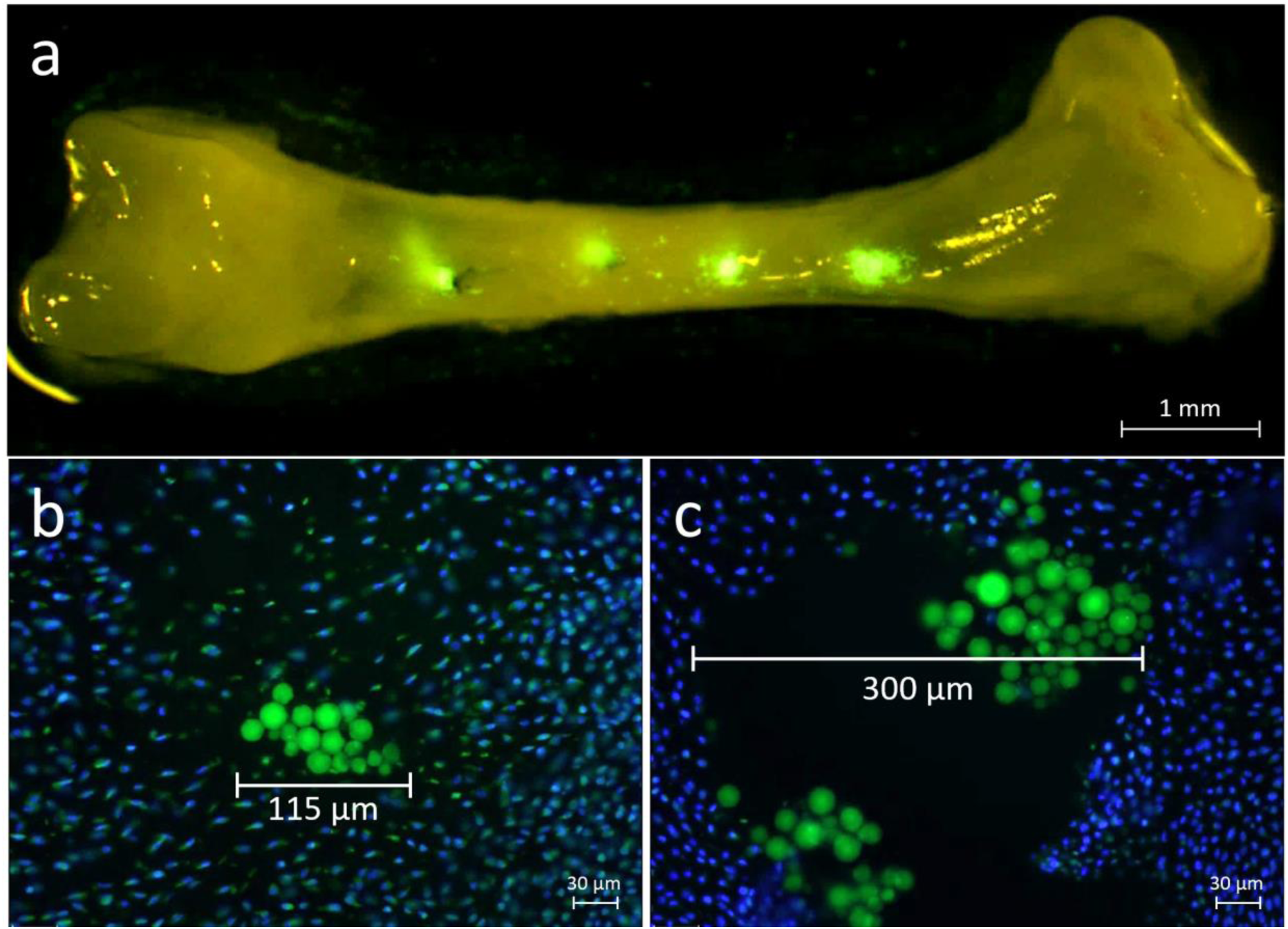
MPs injected in both diaphyseal and epiphyseal regions (a). Minimal invasive damage achieved following microinjection of MPs using glass needle with long and narrow tips with external diameters of ∼100 μm (b). Extensive damage recorded following microinjection using glass needle with shorter tips and wider orifice (c). Nucleus counterstained with 4’, 6-diamidino-2-phenylindole (DAPI).

### 3.2. BMP2-MPs injection

To determine the cellular response to the delivered growth factors following injection, Alcian blue/Sirius red staining was performed on injected samples. To ensure specificity of response, various control groups were included in the study including injection of 1% HSA solution (carrier protein), BMP2 solution and blank PLGA microparticles containing 1%HSA. Microscopical analysis of sections of injected samples showed that morphological changes were limited to reformation of tissue surrounding the injection sites in control samples with no obvious cellular response. However, distinct morphological changes were observed in cells located in close proximity to the BMP2-MPs injection sites in the epiphyseal proliferative zone (Figure 2, PZ). Injection of BMP2-MPs into a zone containing pre-hypertrophic and hypertrophic zones (committed zone - CZ) demonstrated deposition of collagen (collagen fibers evidenced by Sirius red staining) rather than morphological changes (Figure 2).

**Figure 2:**
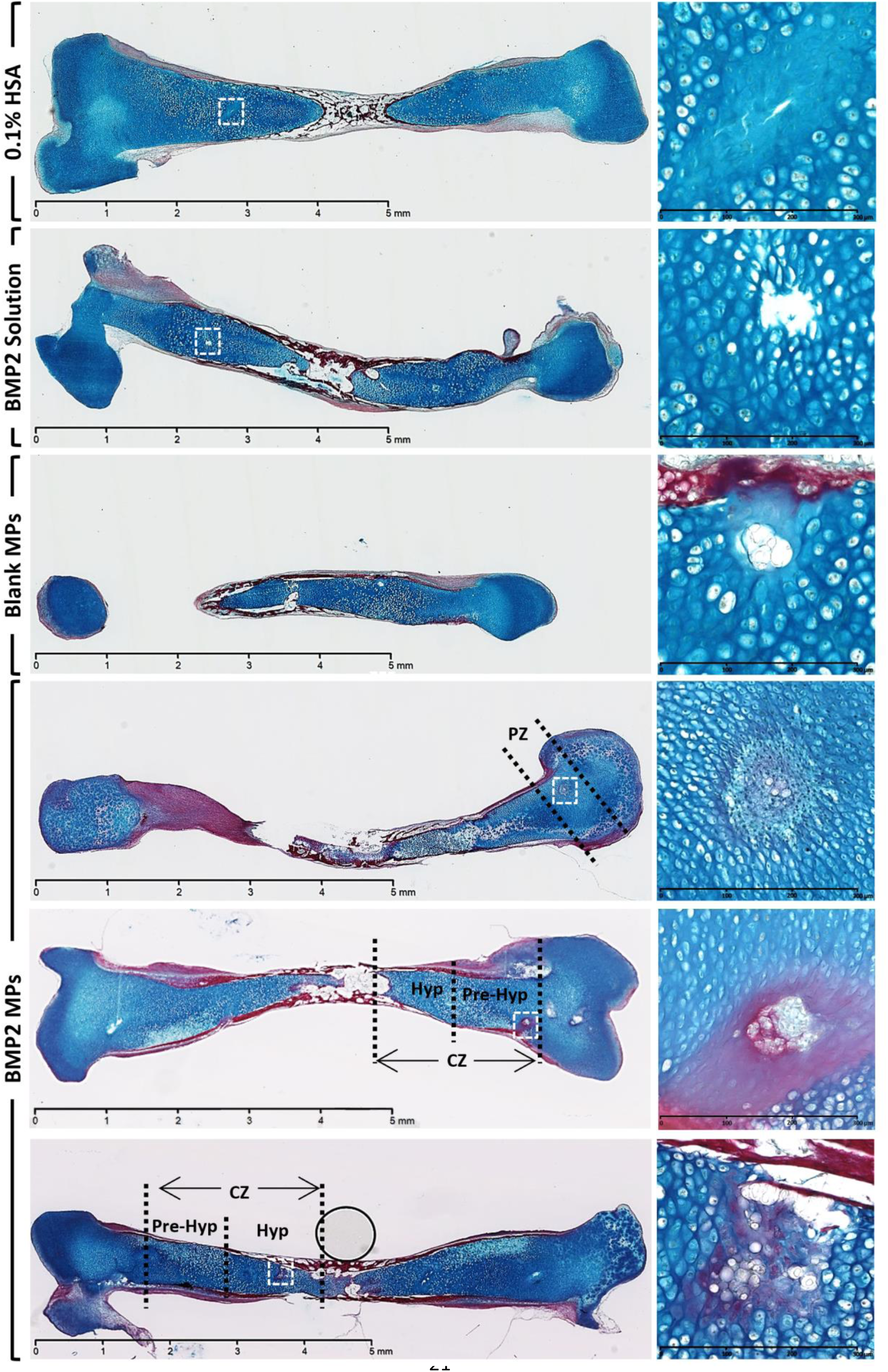
Histological analysis of injected chick bones stained with Alcian blue/Sirius red staining with injection sites highlighted by arrows. Morphological changes in 1% HSA, BMP2 solution and blank MPs controls were subtle and limited to changes in physical disturbance of tissue as a consequence of intervention. In bones injected with BMP2-MPs, morphological changes were observed in a zone of approximately 150 μm in diameter around injection site in proliferative zone (PZ) and deposition of collagen into ECM in samples injected in region contained prehypertrophic (Pre-Hyp) or hypertrophic (Hyp) chondrocytes, here described as committed zone (CZ).

To further analyze the morphology of cells, BMP2-MPs injected samples were analyzed using ESEM. While native cells in the PZ of non-injected samples remained arranged in a dispersed fashion and exhibited the typical round morphology with no distinguishable cytoplasmic projections, cells in close proximity of injected BMP2-MP demonstrated several cytoplasmic projections within a compact arrangement (Figure 3).

**Figure 3:**
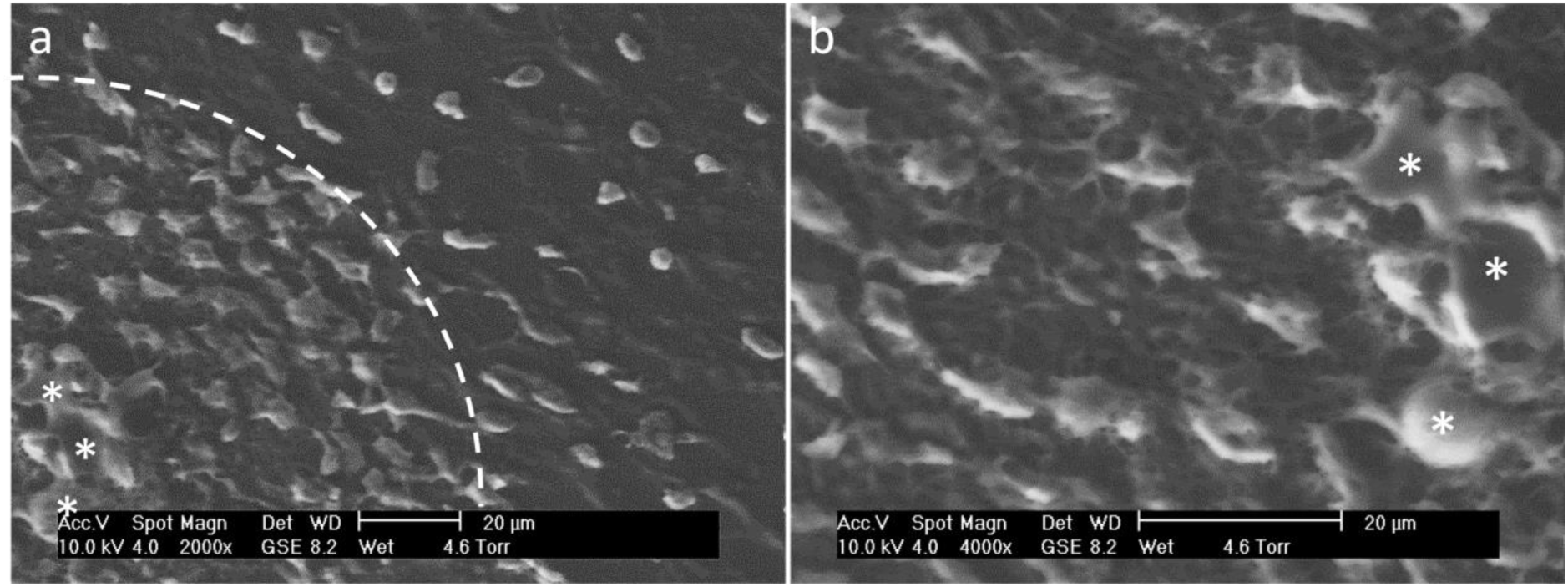
ESEM analysis of injection sites revealed extensive morphological alteration in cells located in close proximity to injected BMP2-MPs (indicated with *). Whilst endogenous cells exhibited round morphology and were located apart from each other (a, outside of the hatched area), cells in close proximity to injected BMP2-MPs arranged in more compact fashion and exhibited an altered morphology with distinct cytoplasmic projections (a, inside hatched area and b).

To examine the expression of osteogenic and chondrogenic markers, IHC analyses of samples were performed using antibodies against COL-I, COL–II, COL-X and OC. Since each of these osteogenic and chondrogenic markers are regulated differentially within the PZ and CZ (based on the level of progenitor commitment), a pronounced difference in level of expression in comparison with neighboring cells was considered as a positive response in injected or control groups. Expression of both COL-I and COL–II was noted to be up-regulated in PZ cells located in close proximity to injected BMP2-MPs in comparison to uncommitted native counterparts. In contrast, injection of BMP2-MPs into the CZ led to a more pronounced expression of COL-I (Figure 4). Similarly, up-regulation of COL-I expression was also observed in cells in close proximity to injected BMP2^ORF^-MPs (Supplementary Figure 4). Differential expression of COL-X was observed between cells located in close proximity to injected BMP2-MPs within the PZ and CZ. Thus, whilst up-regulation of COL-X was negligible in PZ cells, COL-X expression was distinctly up-regulated in CZ cells in close proximity to injected BMP2-MPs in comparison to neighboring counterparts (Figure 4). To evaluate expression of late osteogenic markers, IHC was performed using OC antibody. Despite detection of OC expression in bone collar and hypertrophic chondrocytes, expression of OC was not detected in cells with altered morphology in both PZ and CZ (Figure 4).

**Figure 4:**
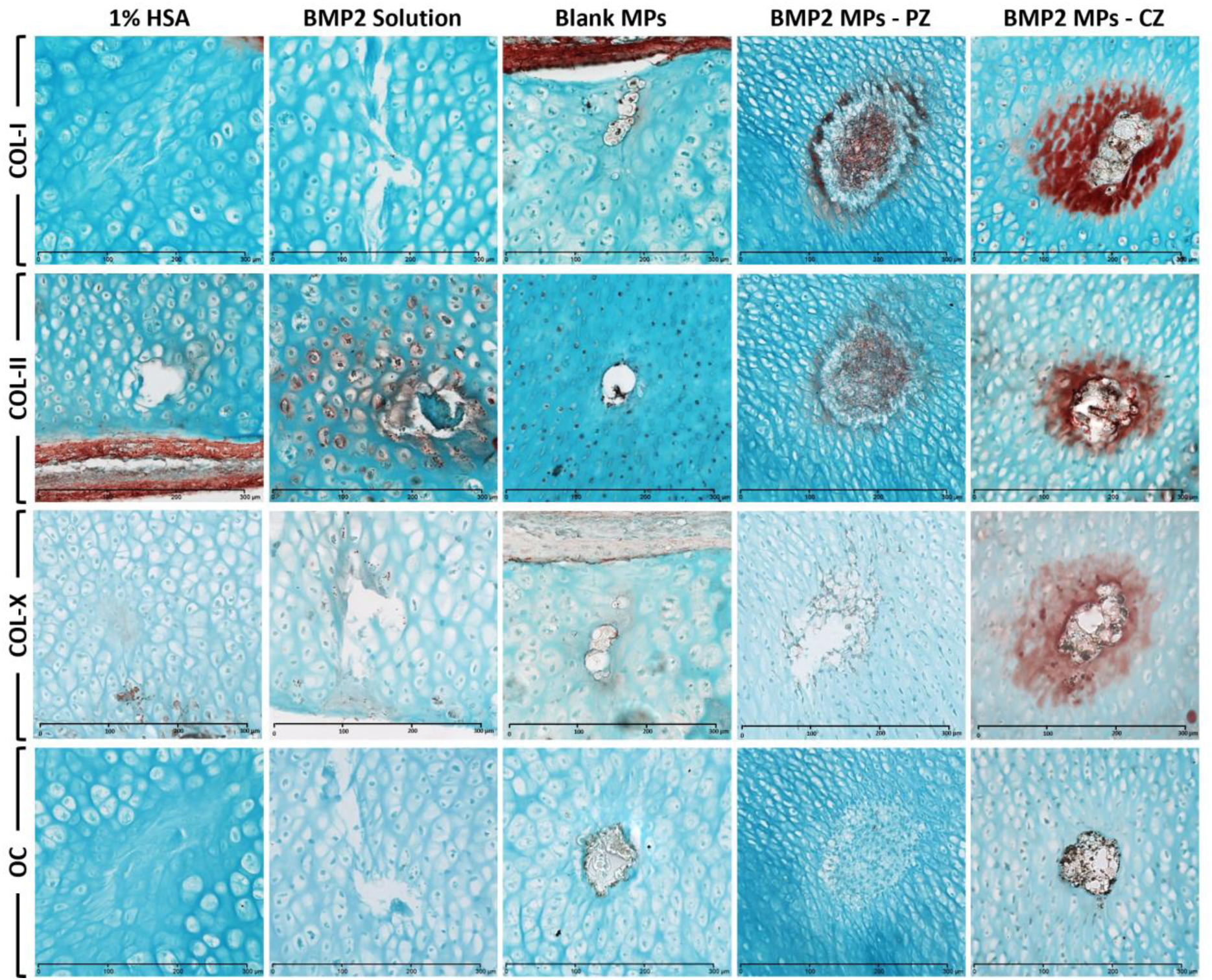
Expression of both COL-I and COL-II was up-regulated in cells around injected BMP2-MP. Expression of COL-I and Col–II (particularly COL-I) was more pronounced in cell located in CZ. Expression of COL-X was negligible in the PZ cells while up-regulation of COL-X was higher in the CZ cells compared to neighboring counterparts. Expression of the late osteogenic marker OC was not detected. No extensive changes were observed in the level of expression of analyzed markers in cells located in close proximity to the injection sites compared with native cells in 1% HSA, BMP2 solution and blank MPS control groups (Please refer to supplementary 5-8 for lower magnification illustration, COL-I, COL-II & OC scale bars: 300 µm, COL-X scale bars: 200 µm).

In analyzed sections from samples with injected BMP2-MPs (n=6), the area of cells displaying morphological changes in chondroprogenitors or the zone of COL-I up-regulation was noted to be clearly limited to proximity of injected MPs covering around 0.0105-0.101 mm^2^ of surface area on each section (Supplementary Figure 5). A macro was developed to measure the intensity of COL-I staining around injected MPs by image segmentation and three level thresholding.

Given the typically low-level COL-I expression in chondroprogenitors, image analysis of samples injected with blank MPs detected only low-intensity, and to a lesser extent, intermediate-intensity signals at the injection sites in the committed zone (CZ), with no detection of the high-intensity segment (Figures 5–6 and Supplementary Figure 6).

**Figure 5.**
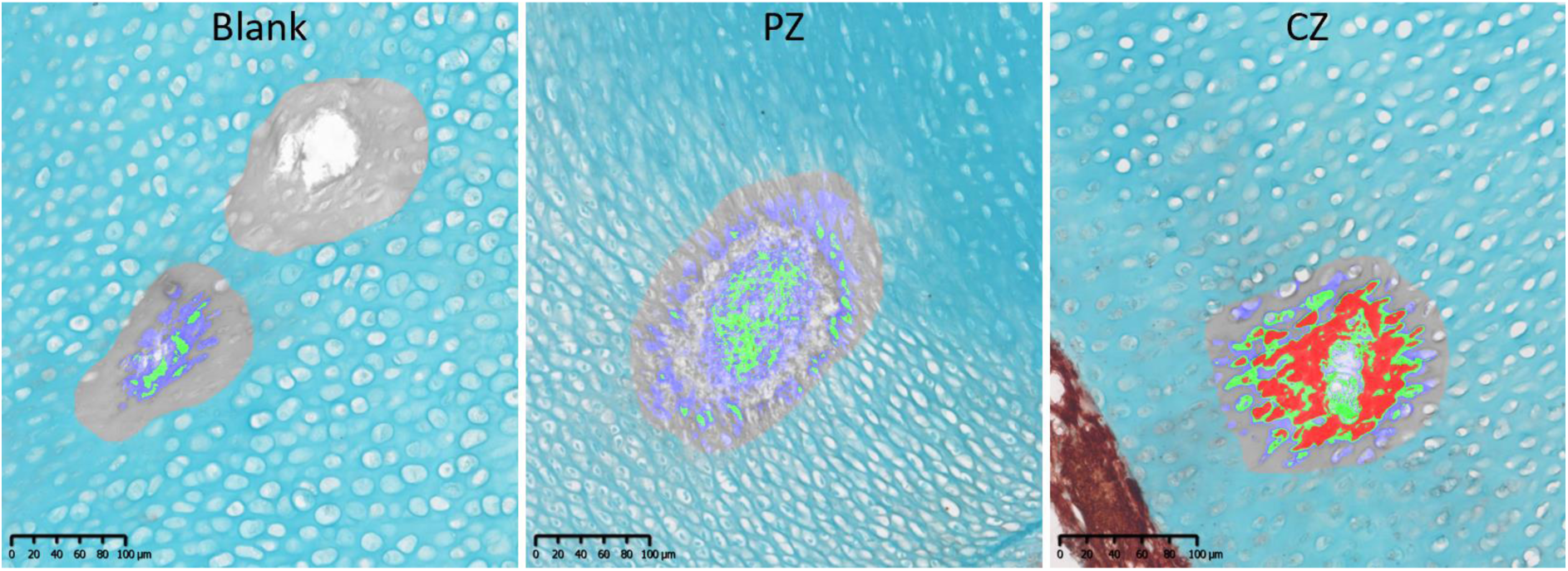
Quantification of COL-I expression following exposure to BMP2 released from GF-loaded MPs based on three-level thresholding where intensity of pixels categorized into >151 (light DAB) as low (blue), 76-150 as intermediate (green) and <75 (dark DAB) as high in chick bone injected with Blank-MPs and BMP2-MPs in proliferative zone (PZ) and pre-hypertrophic zone (CZ).

**Figure 6:**
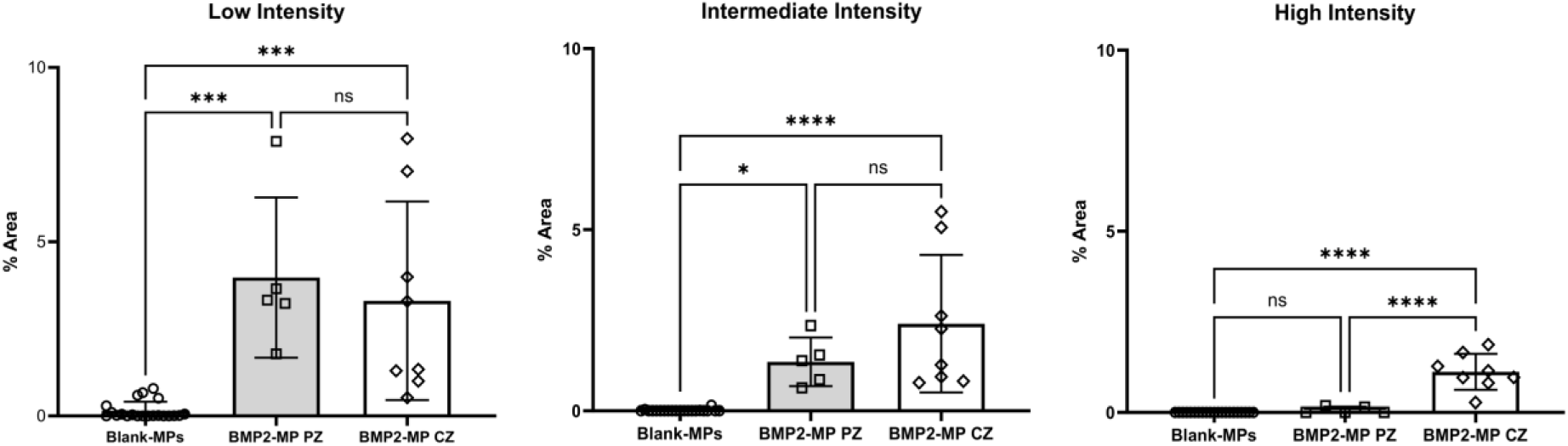
Quantification of COL-I expression area in close vicinity of injection sites. Values represent mean ± SD for Ctrl (n=21), PZ (n=5) and CZ (n=8).

In contrast, sections from samples injected with BMP2-MPs into the PZ exhibited a greater percentage of low- and intermediate-intensity COL-I staining around the injection sites compared to those around blank MPs. Furthermore, areas with high-intensity staining were observed, albeit at a lower proportion relative to the low- and intermediate-intensity signals (Figures 5–6 and Supplementary Figure 7). In CZ, analysis of samples injected with BMP2-MPs revealed a significantly greater percentage of high-intensity COL-I staining compared to those injected in the PZ (Figures 5–6 and Supplementary Figure 8).

### 3.3. FGF2-MPs injection

To evaluate the response of cells located in hyaline cartilage to the spatiotemporal release of FGF2, FGF2-MPs were injected into the chick embryonic bone following a similar approach as adopted for BMP2-MPs. Alcian blue/Sirius red staining of samples following injection of FGF2-MPs revealed that cells located in the PZ and in close proximity to injected FGF2-MPs displayed morphological changes with the formation of extended cytoplasmic projections arranged in random formation (Figure 7). ESEM analysis revealed longer cytoplasmic projections of increased size in PZ cells being exposed to FGF2 which were distinctive from the morphology of cells being exposed to BMP2-MPs (Figure 7). Further analysis of samples revealed differential regulation of COL-I and COL–II in chondrocytes located in both PZ and CZ in comparison with bones injected in the same area with BMP2-MP. While COL-II expression was highly up-regulated in comparison to native counterparts, expression of COL-I was not detected in cells exposed to FGF2-MPs. Similarly, up-regulation of only COL-II was observed in the CZ cells in close proximity to injected FGF2-MPs. A low level of COL-X expression was observed in CZ cells located around the injection site (Figure 8).

**Figure 7:**
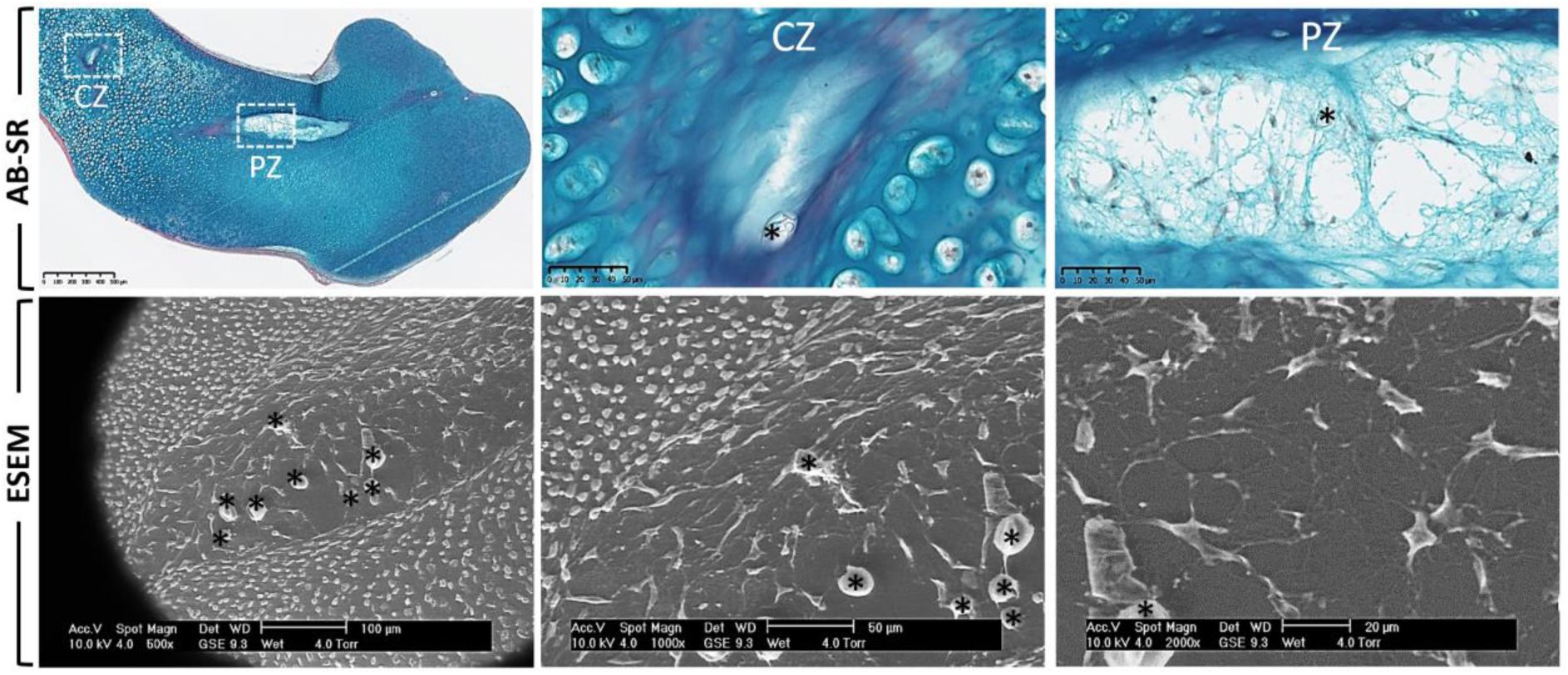
Morphological changes was observed following transplantation of FGF2-MPs in sections stained with Alcian Blue/Sirius Red (AB-SR) which was more distinctive in PZ cells. The appearance and arrangement of morphologically changed cells were distinctive from PZ cells in close proximity to BMP2-MPs. No obvious alteration was observed in morphology of cells located in CZs. Following ESEM, longer cytoplasmic projections were observed in cells located in PZ following exposure to sustained release of FGF2 from MPs (indicated with *) and they re-arranged in a more dispersed fashion. These cells displayed a distinct morphology compared to cells exposed to BMP2.

**Figure 8:**
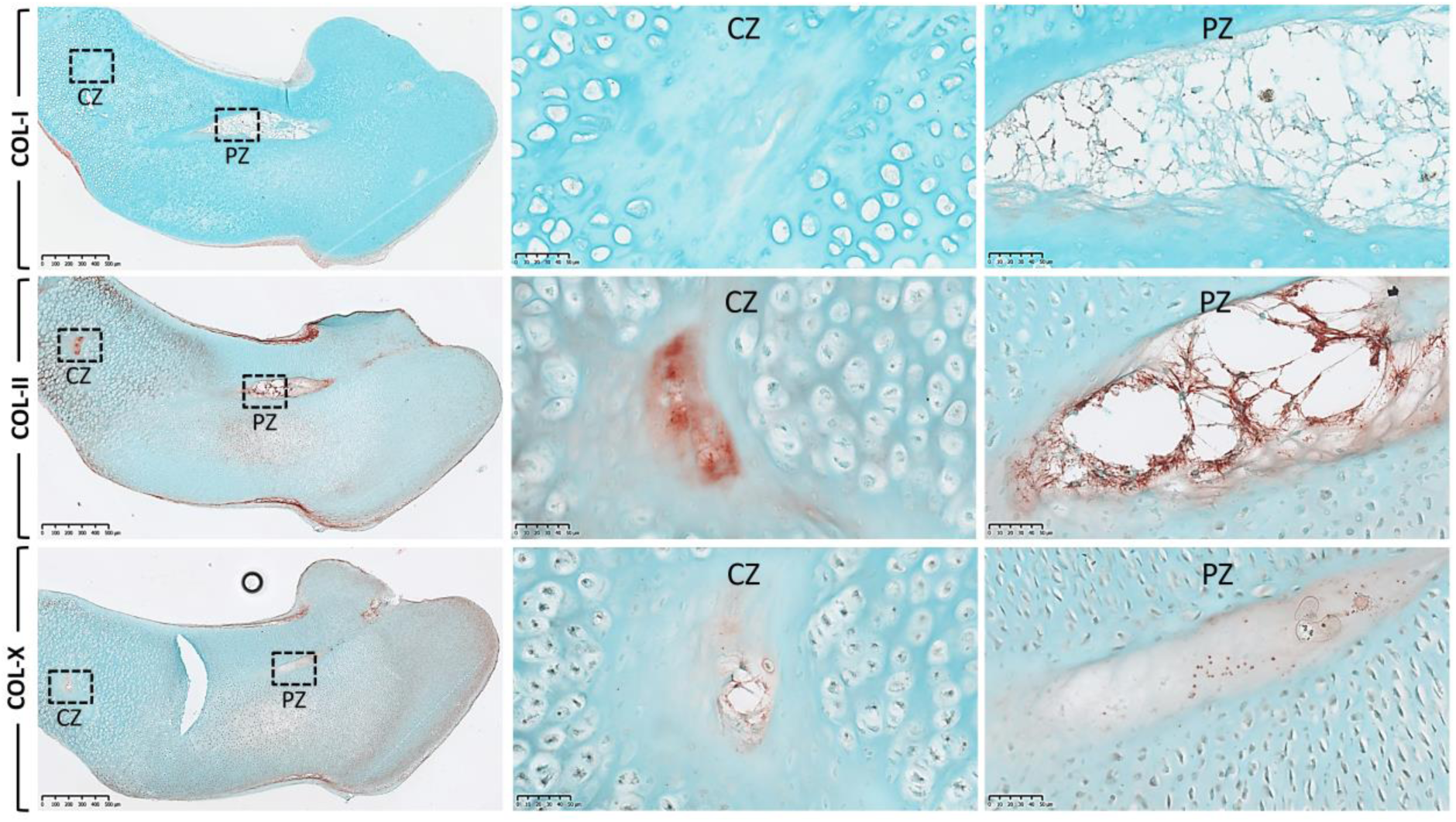
Up-regulation of COL-II was observed in chondroprogenitors in the PZ and CZ in closed proximity to the injection sites. No distinctive changes were observed in expression of COL-I and COL-X in cells located in close proximity to the injected FGF2-MPs compare with native cells.

## 4. Discussion

In this study we have highlighted the significant effect that spatio-temporal regulation of growth factor signaling has on tissue formation. We used PLGA microparticles to deliver and controllably release BMP2 and FGF2 in an *ex vivo* embryonic chick femur, which was used as a model of bone formation.

The noted expression of COL-I in response to the microparticle delivered BMP2 indicated osteogenic commitment of chondroprogenitors and potential early onset of mineralization of matrix as described previously (Owen et al., 1990). In contrast, the lack of OC expression, suggests limited osteoblast differentiation and maturation (Lynch et al., 1995). The lack of expression of OC and evidenced osteoblast maturation of skeletal populations may be a consequence of various factors including short exposure time and or constant exposure to BMP2, which has been previously reported to delay terminal differentiation of hypertrophic chondrocytes (Minina et al., 2001, Minina et al., 2002). In addition, inhibitory signals released from adjacent chondroprogenitors such as tenascin-w may play role and inhibit further differentiation of exposed cells (Kimura et al., 2007). The observation that injection of BMP2 solution had a negligible effect on cells in close proximity to injection sites further substantiates the importance of a controlled-release strategy to induce the desired clinical effect and to improve reparative outcomes. As bacterial endotoxin has been shown to elicit cellular responses (Magnusdottir et al., 2013), endotoxin-free BMP2-loaded MPs (BMP2^ORF^-MPs) were injected following a similar procedure to rule out the potential effects of endotoxin. Up-regulation of COL-I in cells in close proximity to injected BMP2^ORF^-MPs suggested that the observed cellular response is a direct consequence of exposure to BMP2 released from MPs rather than bacterial endotoxin (Supplementary 4).

Fibroblast growth factors (FGFs) are important regulators of osteogenesis while their importance in bone tissue engineering and, critically, their precise role remains far from clear. Increased cellular proliferation and enhanced adipogenic differentiation (Neubauer et al., 2004) have been reported as well as inhibition of osteogenic differentiation (Quarto and Longaker, 2006) following exposure to FGF2. Interestingly, synergistic effects of FGF2 and BMP2 on osteogenic differentiation of bone marrow-derived mesenchymal stem cells have been reported elsewhere (Bai et al., 2013, Kuhn et al., 2013). Furthermore, Chiou and colleagues have shown FGF2 has mitogenic and chondrogenic effects on adipose-derived MSCs (Chiou et al., 2006). The noted lack of COL-I expression and up-regulation of COL-II following exposure to FGF2 in the current studies is rather in accordance with previously published data suggesting FGF2 as a negative regulator of osteoblast differentiation and enhancer of chondrogenesis (Quarto and Longaker, 2006, Chiou et al., 2006). Moreover, the distinct regulation of COL-I and –II in cells located in close proximity to the injected BMP2-MPs in contrast to FGF2-MPs suggests a growth factor-dependent response to exogenous factors. These observations further emphasize the importance of a physiologically-relevant environment to delineate the effect of extrinsic inductive factors.

Previously, a modified *ex vivo* chick embryonic femur culture system was developed to study the effect of osteoinductive factors (Kanczler et al., 2012, Smith et al., 2015). In addition, this organotypic model has been used as a critical bone defect model to study the effect of growth factor released from microparticles encapsulated in an alginate construct and implanted in a bone defect site created in the bone diaphysis (Smith et al., 2014, Gothard et al., 2024).

To study the localized effect of spatiotemporal release of GFs, here we introduce a new approach by taking advantage of a recently developed and modified *ex vivo* chick embryonic femur culture system (Kanczler et al., 2012). The lack of response in control groups injected with BMP2 solution indicates the importance of an efficient sustained release mechanism to fully harness GF potential. In the current study, growth factor-dependent responses of chondroprogenitors indicated the usefulness of this *ex vivo* system in evaluating the effect of GF released from MPs on tissue formation. In addition, this model has the potential to be used for the evaluation of more complex regimens based on the delivery of multiple growth factors. Therapeutic efficacy can be achieved, to a certain degree, following the delivery of a single exogenous factor, physiological systems typically necessitate a cascade of factors and the need for delivery platforms capable of providing spatiotemporal release of several, ideally inductive, factors to mimic aspects of the natural healing process (Mehta et al., 2012). To achieve such a level of sophistication and to evaluate the outcome, approaches have included high throughput screening (Anderson et al., 2005), organ on a chip or microfluidic 3D cell culture devices (Baker, 2011, Polini et al., 2014), and mathematical modelling (Kanjickal and Lopina, 2004, Tzafriri et al., 2005).

## 5. Conclusion

The routine treatment of critical-sized bone defects remains a challenging unmet need (Roddy et al., 2018). Enhanced understanding of the interaction of skeletal progenitor cells with chondro-, osteo-, and angiogenic factors is pivotal to the development of innovative efficacious strategies to mimic endochondral ossification using bone tissue engineering approaches (Owen and Reilly, 2018). However, current *in vitro* assays are limited in their ability to provide all the requisite information.

Our results suggest that the organotypic chick embryonic bone provides a useful *ex vivo* tool to evaluate the activity of multiple growth factors within more physiologically relevant niches. This innovative approach offers much potential as an inexpensive physiologically relevant *ex vivo* model to bridge the gap between current *in vitro* assays and challenging and expensive *in vivo* critical-sized defect animal models.

## Acknowledgment

This work was supported by the strategic longer and larger grant (sLOLA) from the Biotechnology and Biological Sciences Research Council (BBSRC), UK-grant number BB/ G010579/1. We gratefully acknowledge Mrs Nicola Weston at University of Nottingham for ESEM imaging. All research at Great Ormond Street Hospital NHS Foundation Trust and UCL Great Ormond Street Institute of Child Health is made possible by the NIHR Great Ormond Street Hospital Biomedical Research Centre. The views expressed are those of the author(s) and not necessarily those of the NHS, the NIHR or the Department of Health. For the purpose of open access, the author has applied a CC BY public copyright license to any Author Accepted Manuscript version arising from this submission.

## Appendix A. Supplementary data

**Supplementary Figure 1:**
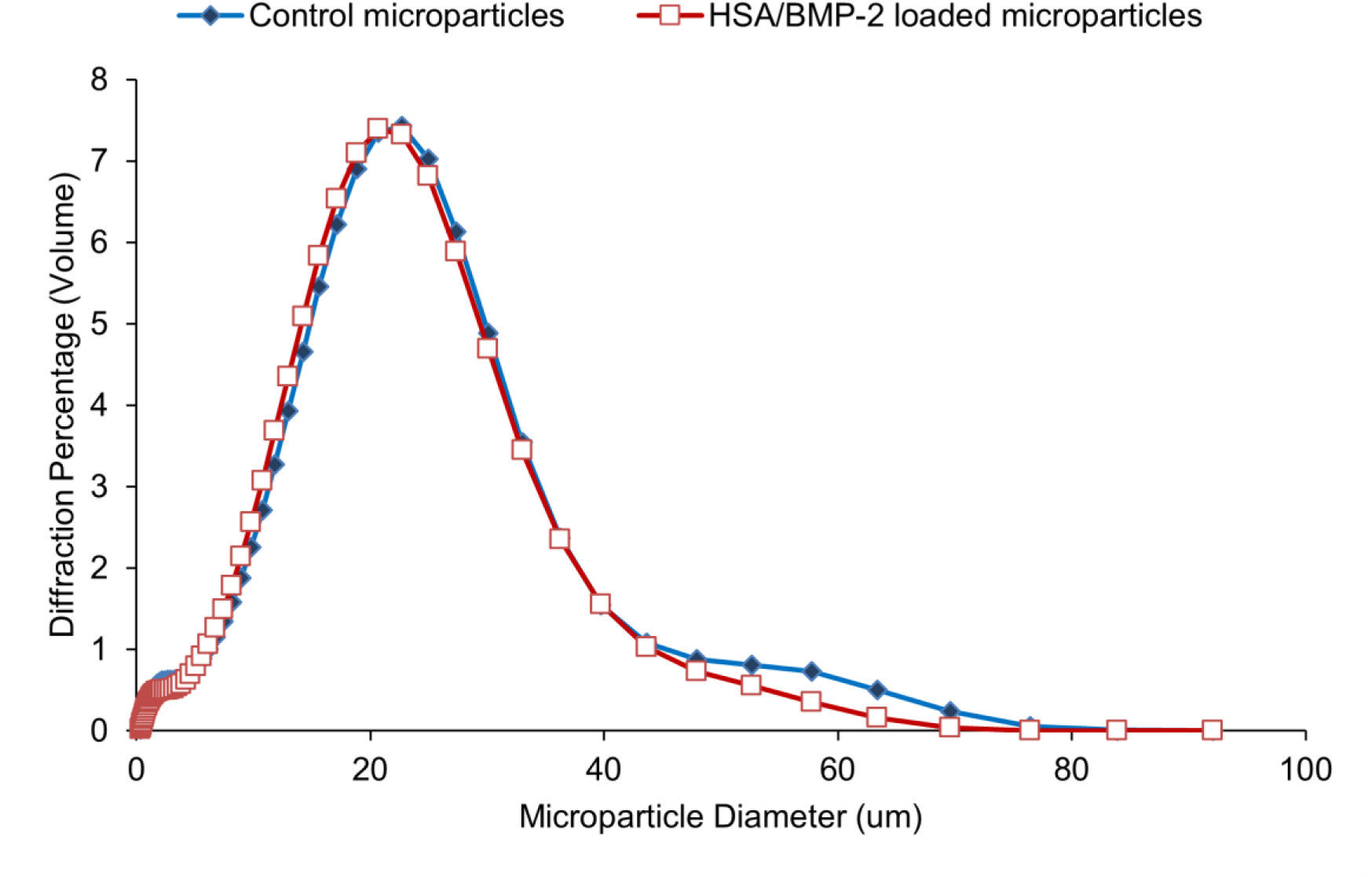
Size distribution of HSA/BMP-2 (1% w/w) loaded PLGA 50:50 (59KDa) microparticles modified with 10% w/w PLGA-PEG-PLGA triblockk copolymer and comparison with control (no HSA/BMP-2). Results quantified by laser diffraction (Coulter LS230).

**Supplementary Figure 2:**
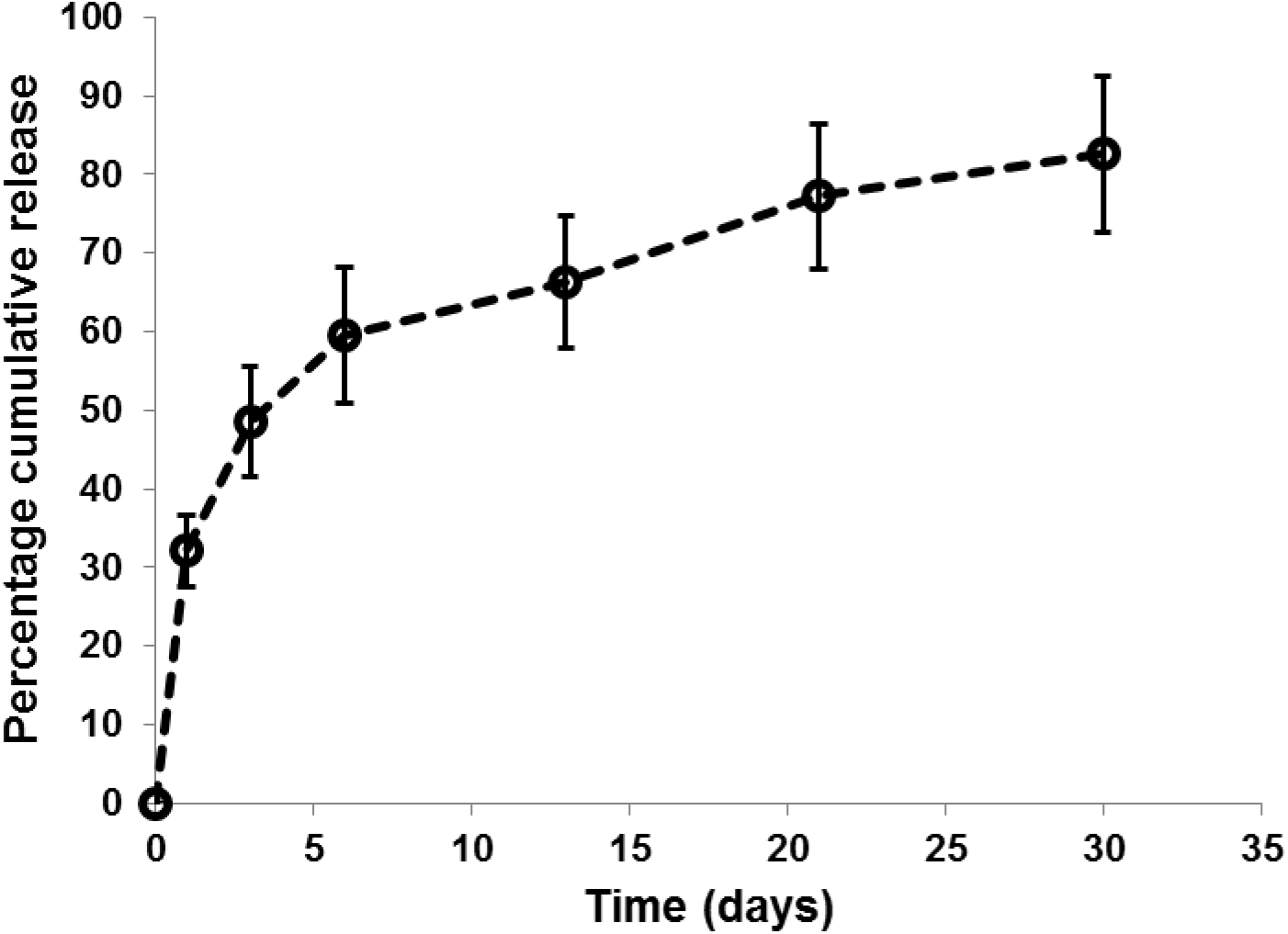
Cumulative *in vitro* release profile of HSA/BMP-2 from PLGA50:50 (59 KDa) microspheres modified with 10% w/w PLGA-PEG-PLGA triblock copolymer manufactured by double emulsion to be 10-40 µm in size. Total protein loading was 10 mg (9.5 mg HSA/ 0.5 mg BMP-2) in 1 g PLGA and release was quantified using a micro-BCA assay.

**Supplementary Figure 3:**
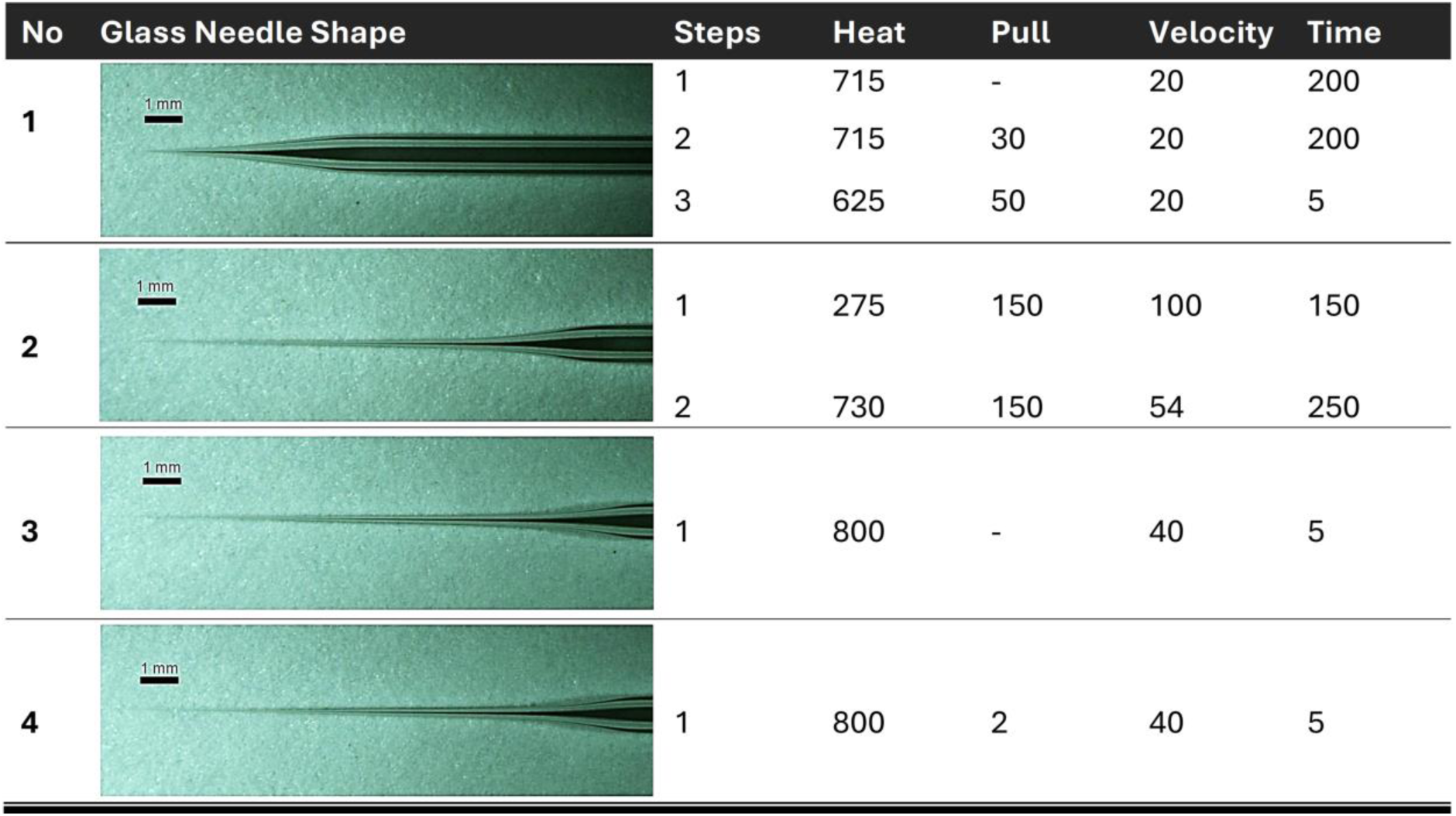
Several glass needles were pulled and tested for optimization of minimal invasive injection of MPs into embryonic chick femur. Minimal invasion was achieved using glass needles no 4 with long tip and an average diameter of ∼50 μm, which was throughout this study.

**Supplementary Figure 4:**
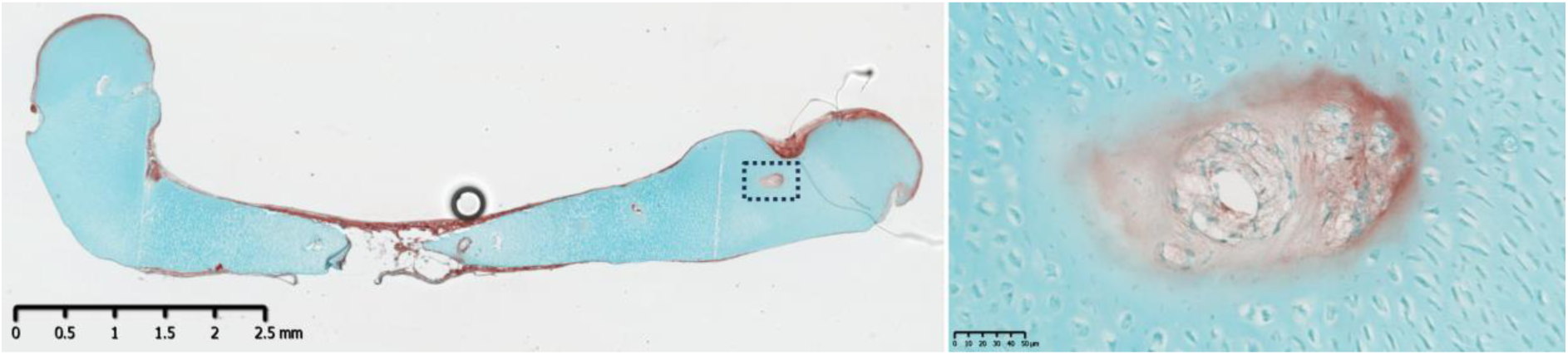
Morphological alteration of chondroprogenitors and up-regulation of COL-I following injection of MPs encapsulated with endotoxin free-BMP2 (BMP2^ORF^-MPs) suggesting that the cellular response is a consequence of BMP2 rather than bacterial endotoxin.

**Supplementary Figure 5:**
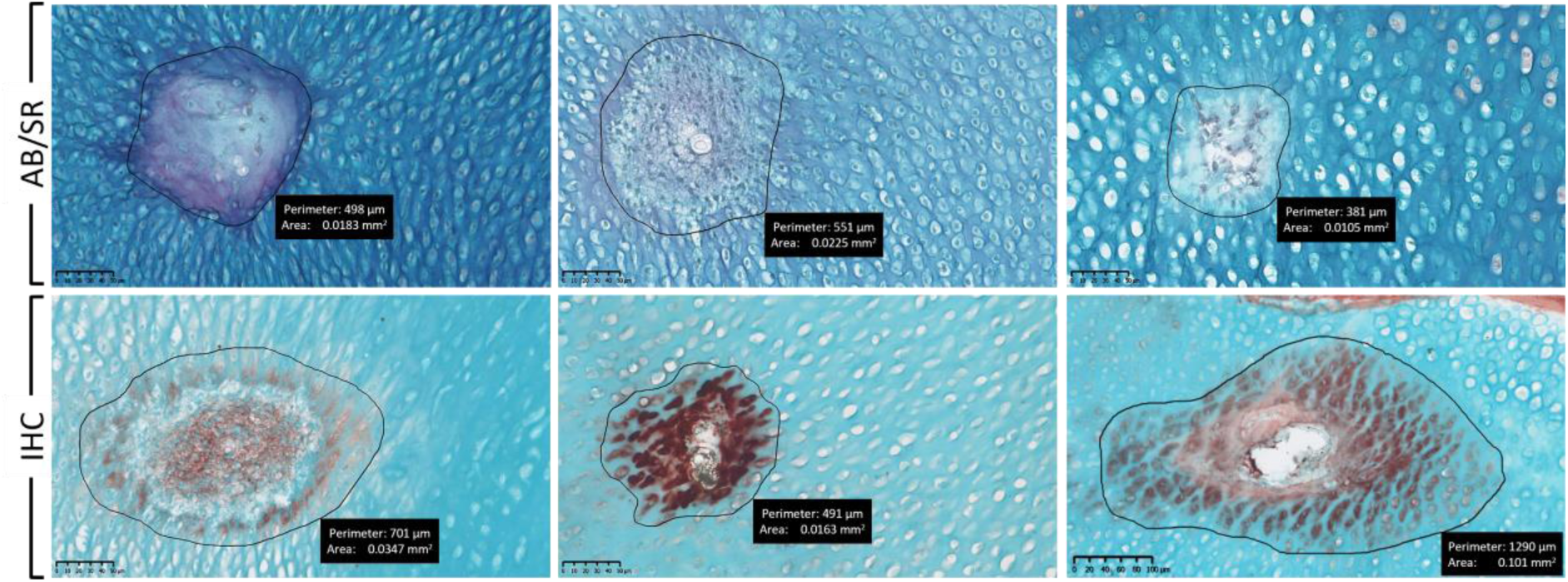
The effect of released BMP2 from MPs on cell morphology and ECM composition (AB/SR) and expression of COL-I as an osteogenic marker (IHC) was localized and limited to close proximity of injected MPs and covered an area of around 0.0105-0.101 mm^2^ in each section.

**Supplementary Figure 6:**
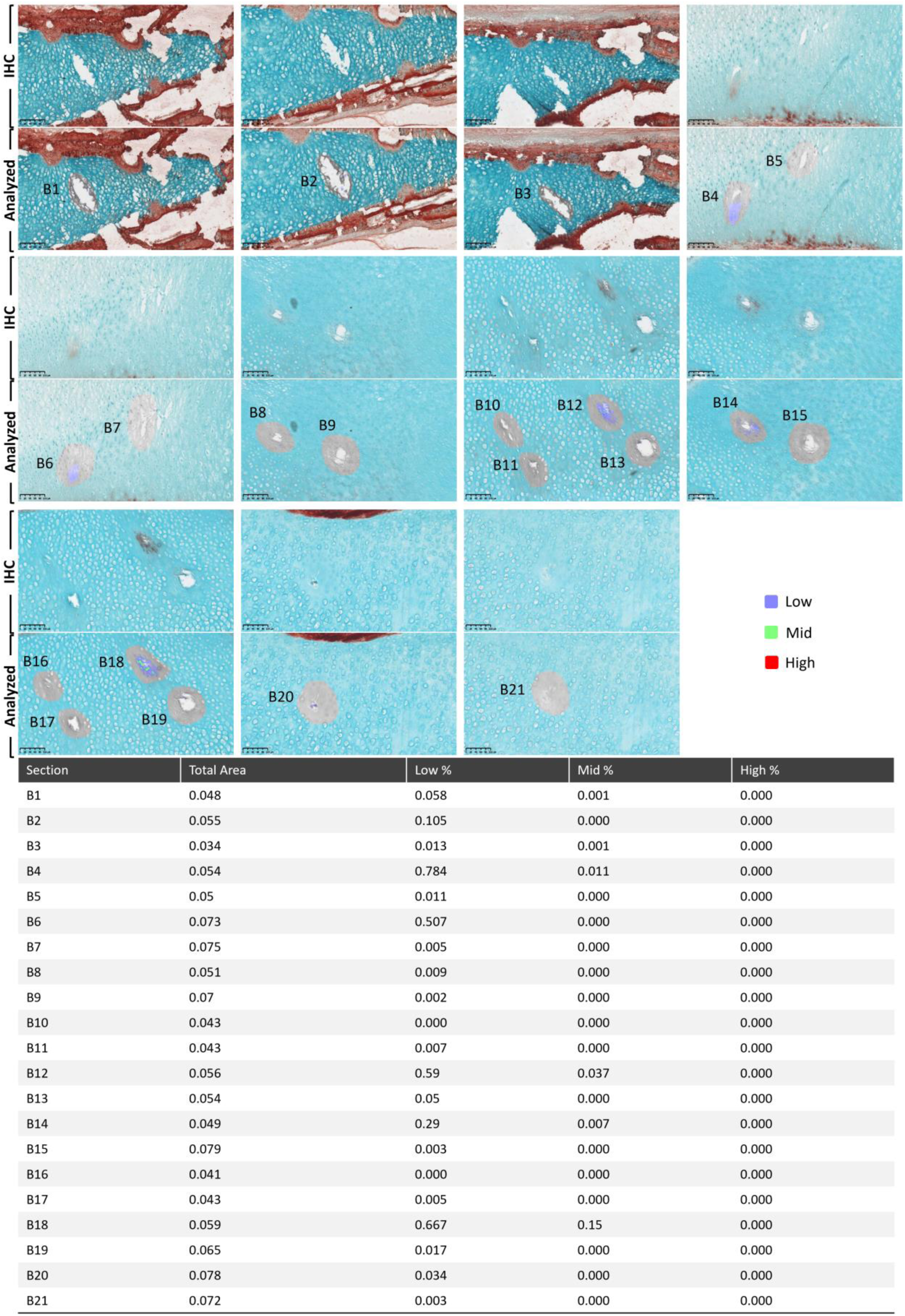
Image segmentation and three-level thresholding of blank-MP injected chick bones. Only areas with low- and intermediate (Mid)-level signals were detected (Scale bars 100 µm).

**Supplementary Figure 7:**
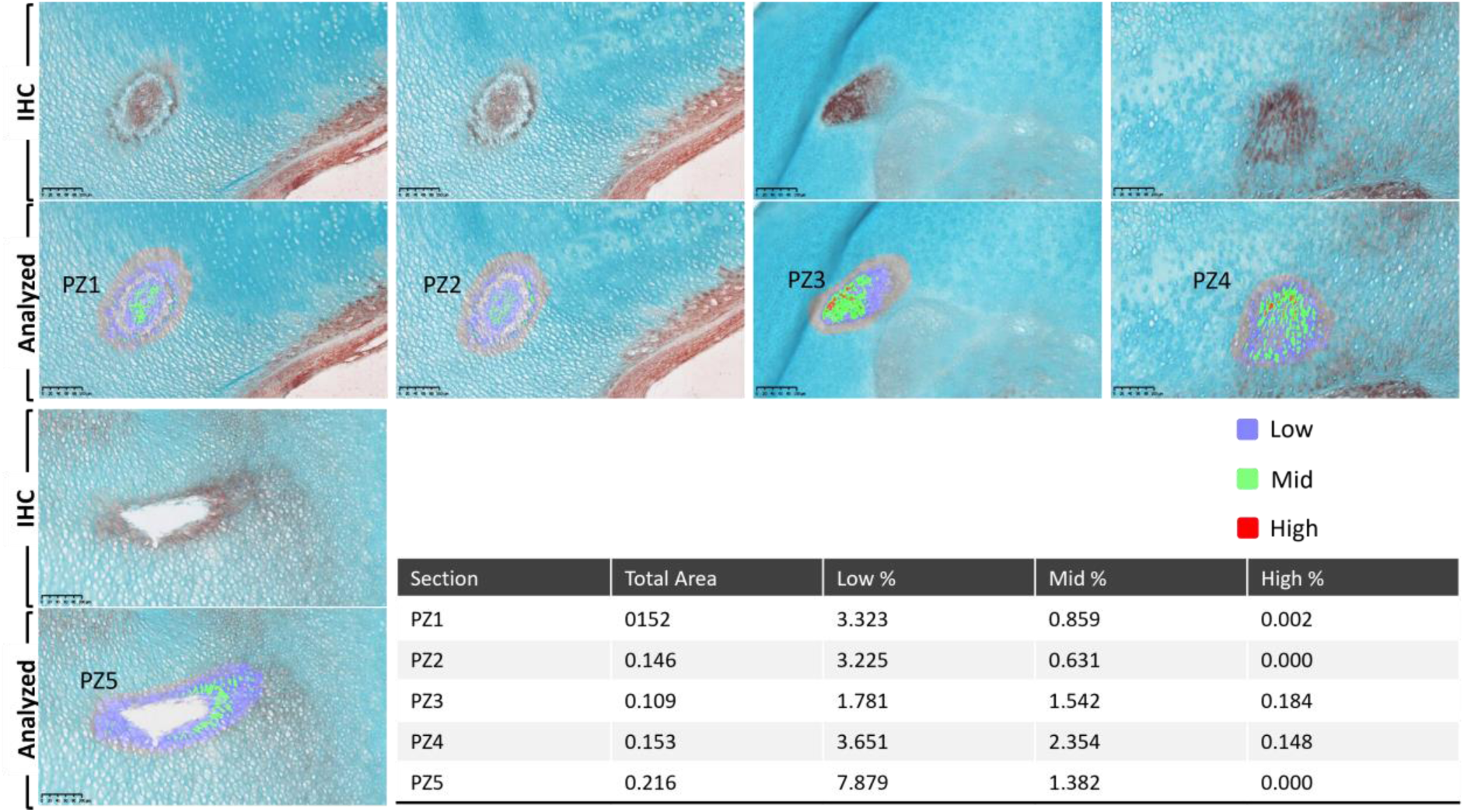
Image segmentation and three-level thresholding of BMP2-MP injected to PZ of chick bones. Higher percentage of areas with low- and intermediate (Mid)-level signals were detected compared to Blank-MP injected samples. Areas with high intensity signal were also detected at lower proportion to areas with low and intermediate level signals (Scale bars 100 µm).

**Supplementary Figure 8:**
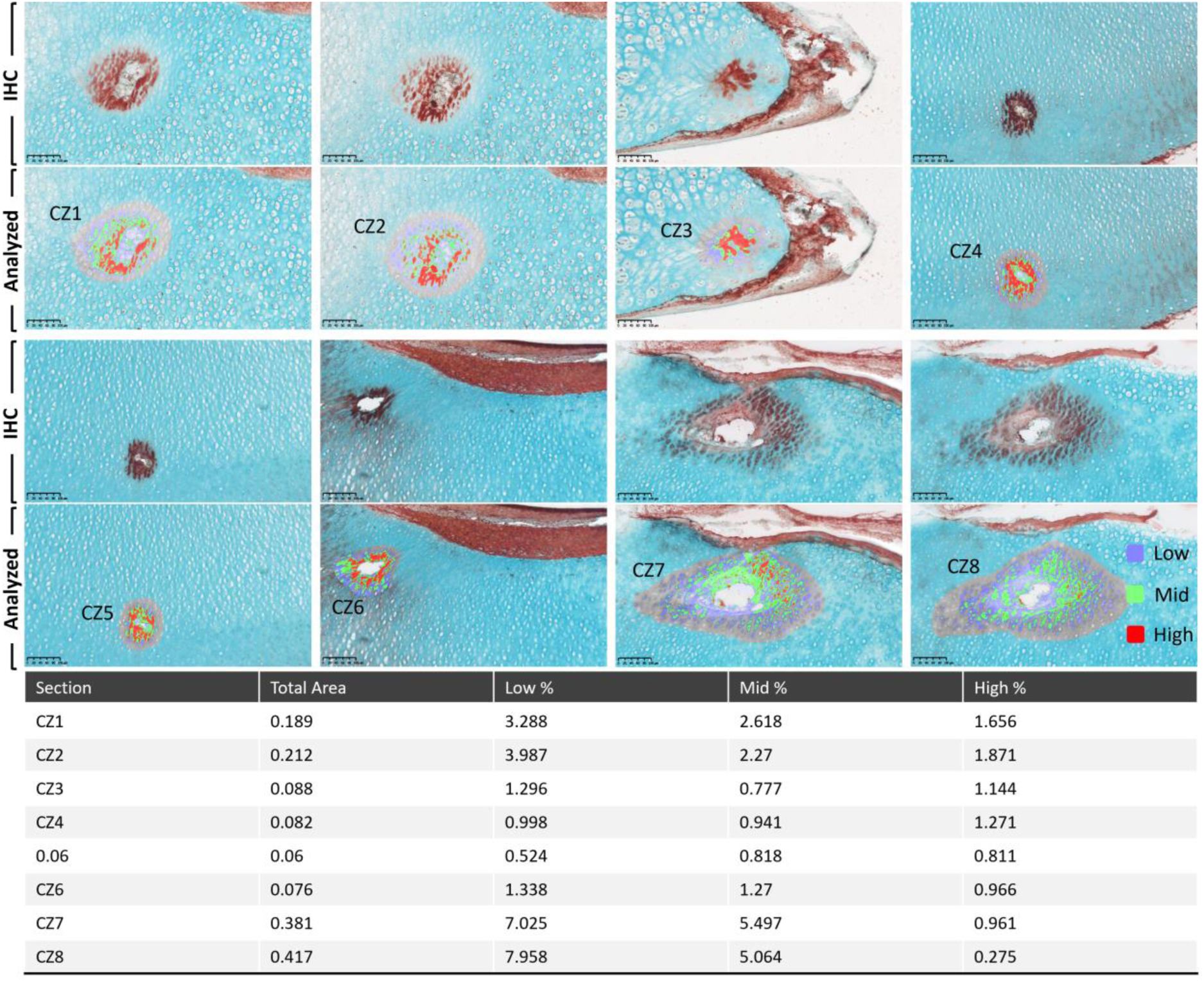
Areas with low-, intermediate (Mid)- and high-level signals were detected following image segmentation and three-level thresholding of BMP2-MPs injected to CZ of chick bones. Proportion of high-intensity areas significantly increased compared to PZ (Scale bars 100 µm).

