## Supplementary figures and images for "Localized Delivery of Growth Factors from Microparticles Modulate Osteogenic and Chondrogenic Gene Expression in Growth Factor-dependent Manner in an *ex vivo* Chick Embryonic Bone Model"

### Supplementary Figure 1

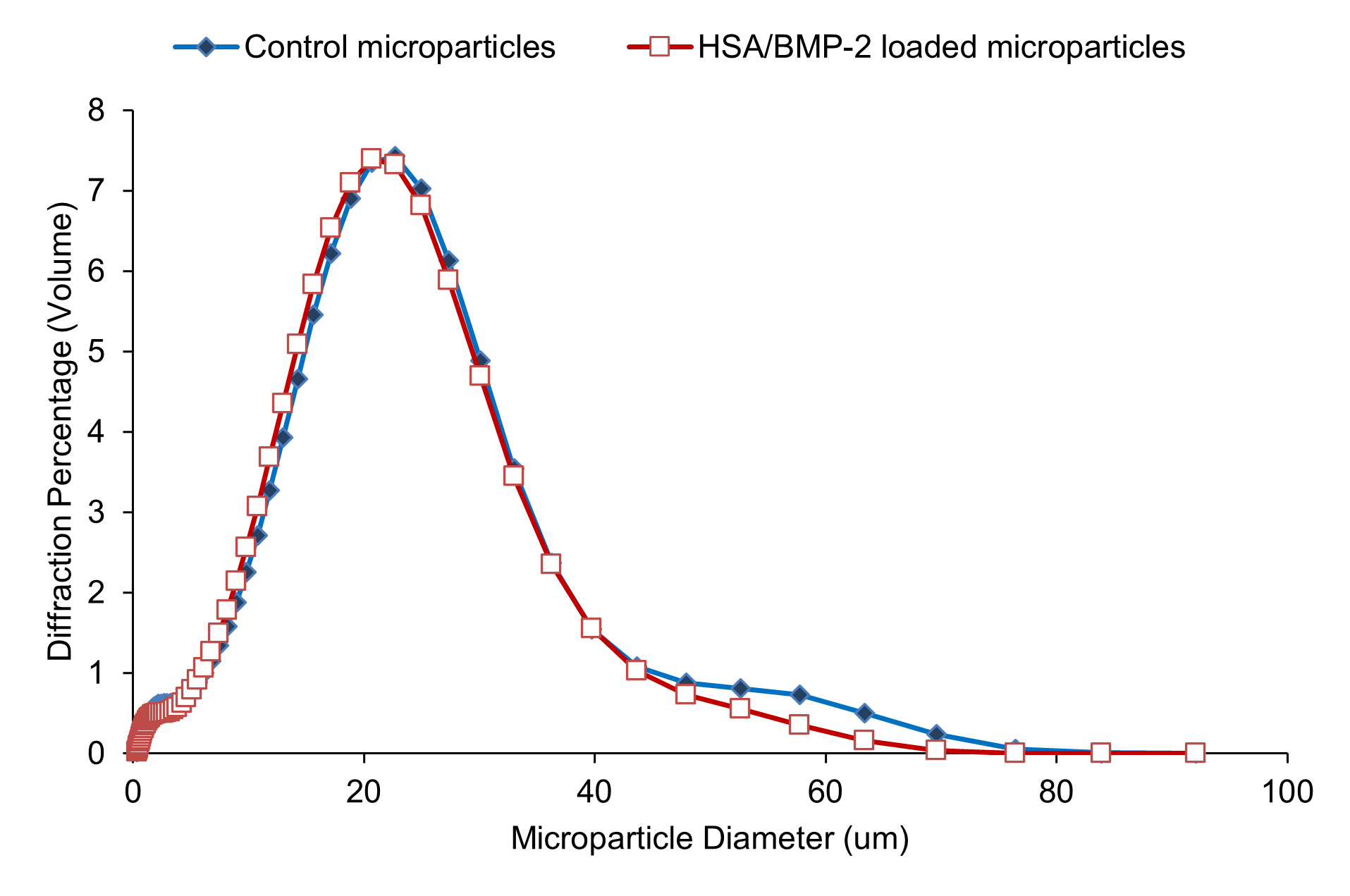

### Supplementary Figure 2

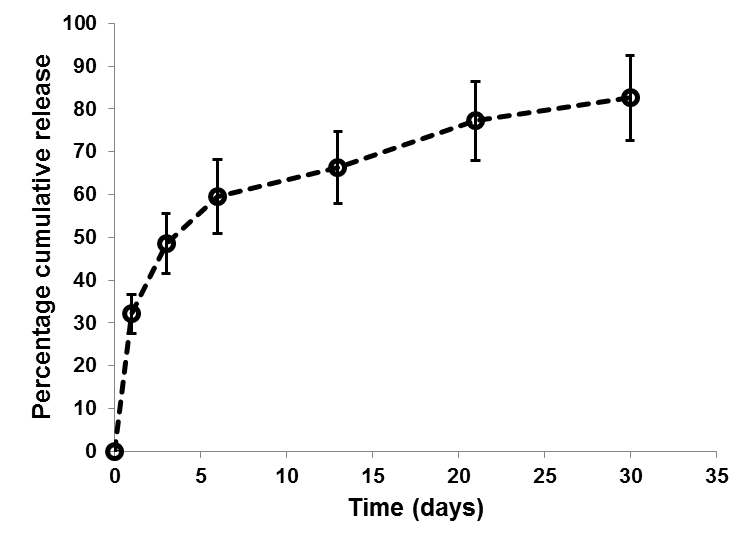

### Supplementary Figure 3

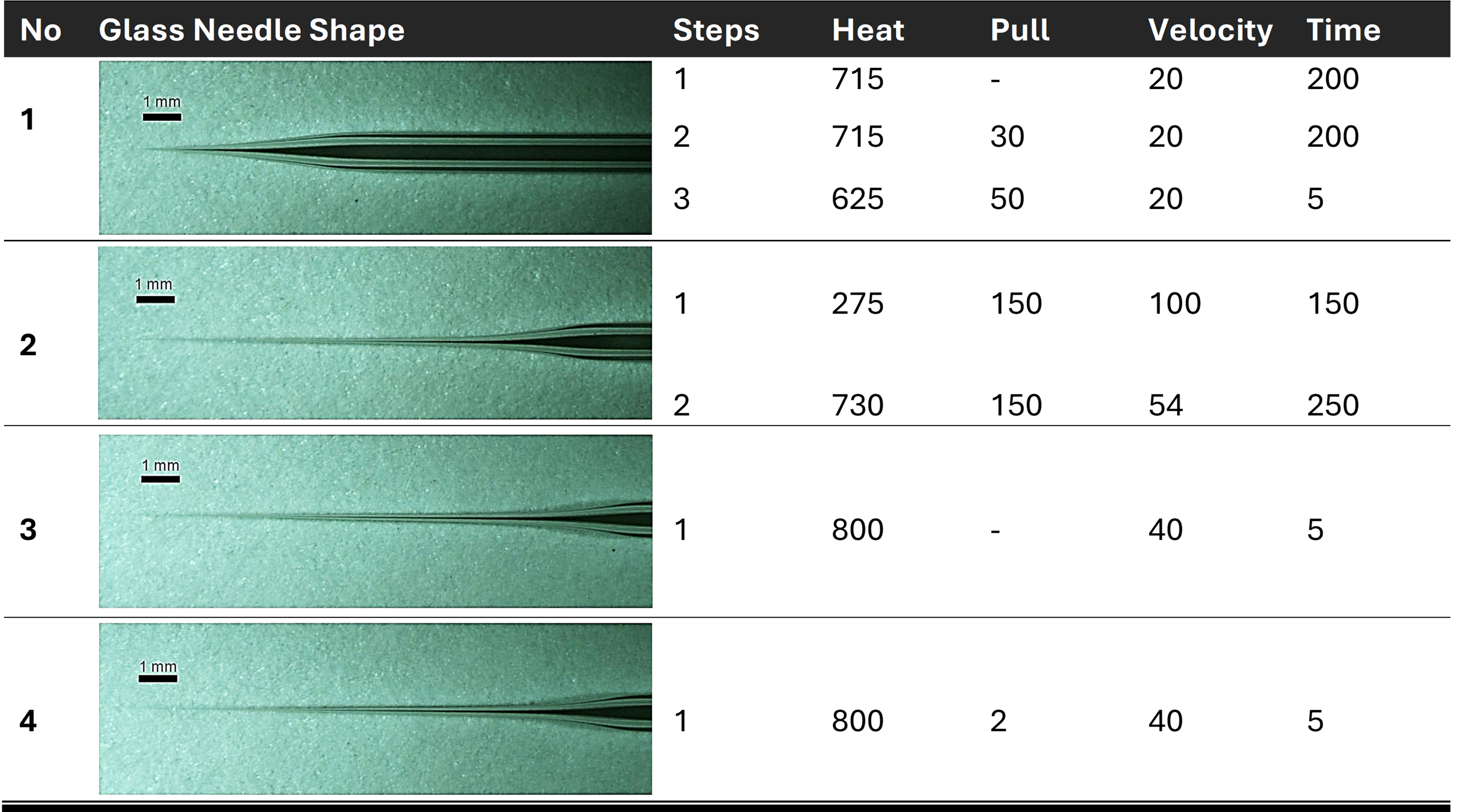

### Supplementary Figure 4

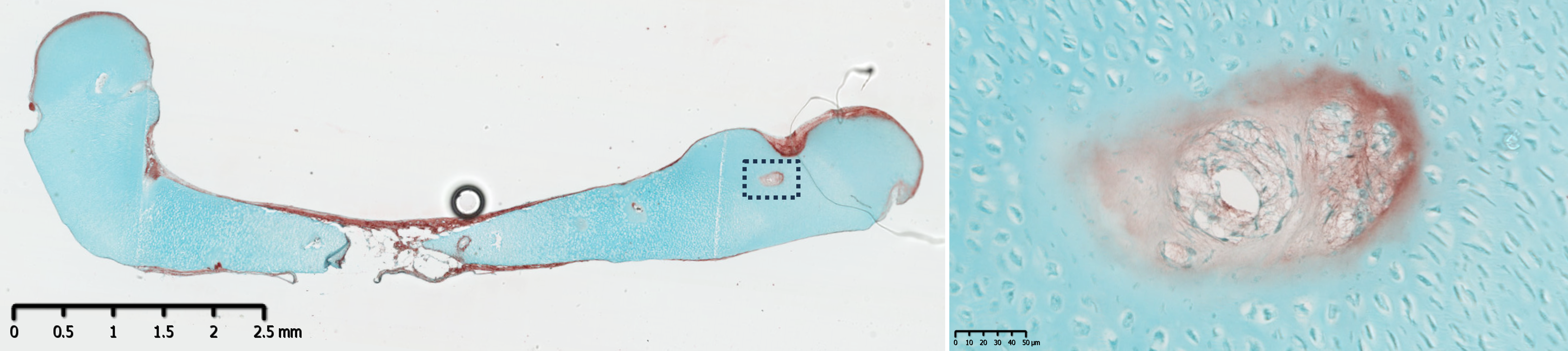

### Supplementary Figure 5

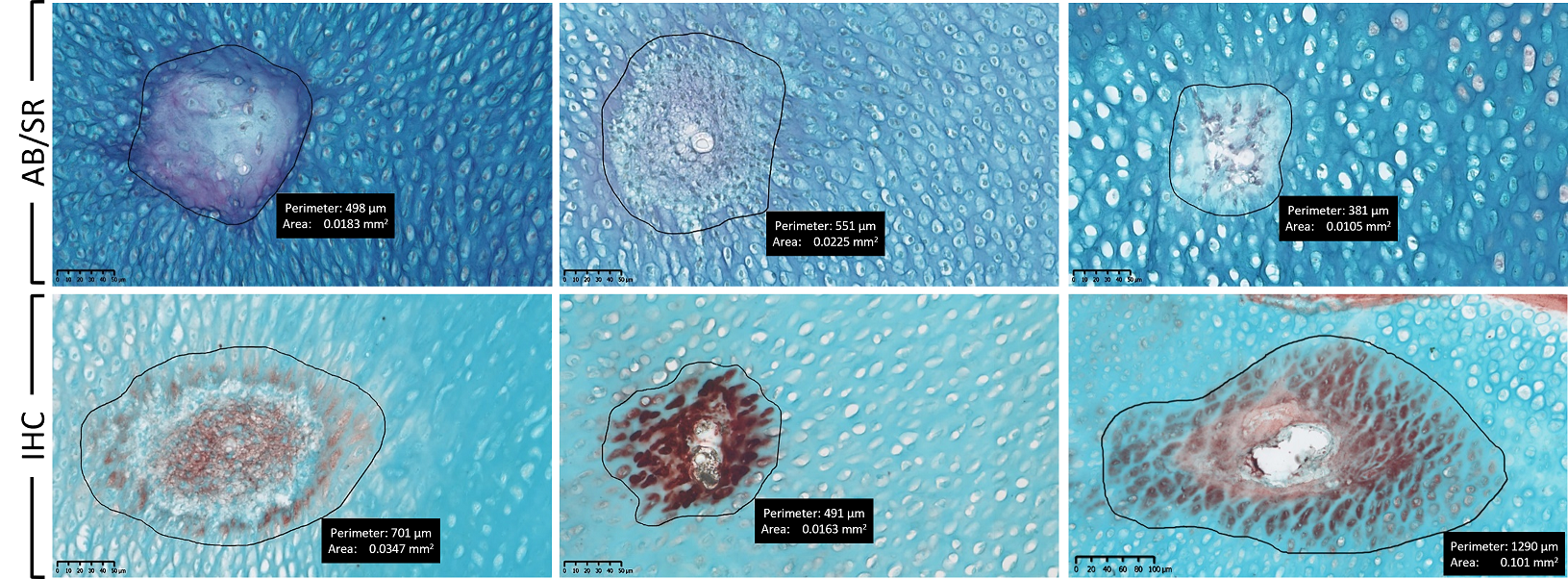

### Supplementary Figure 6

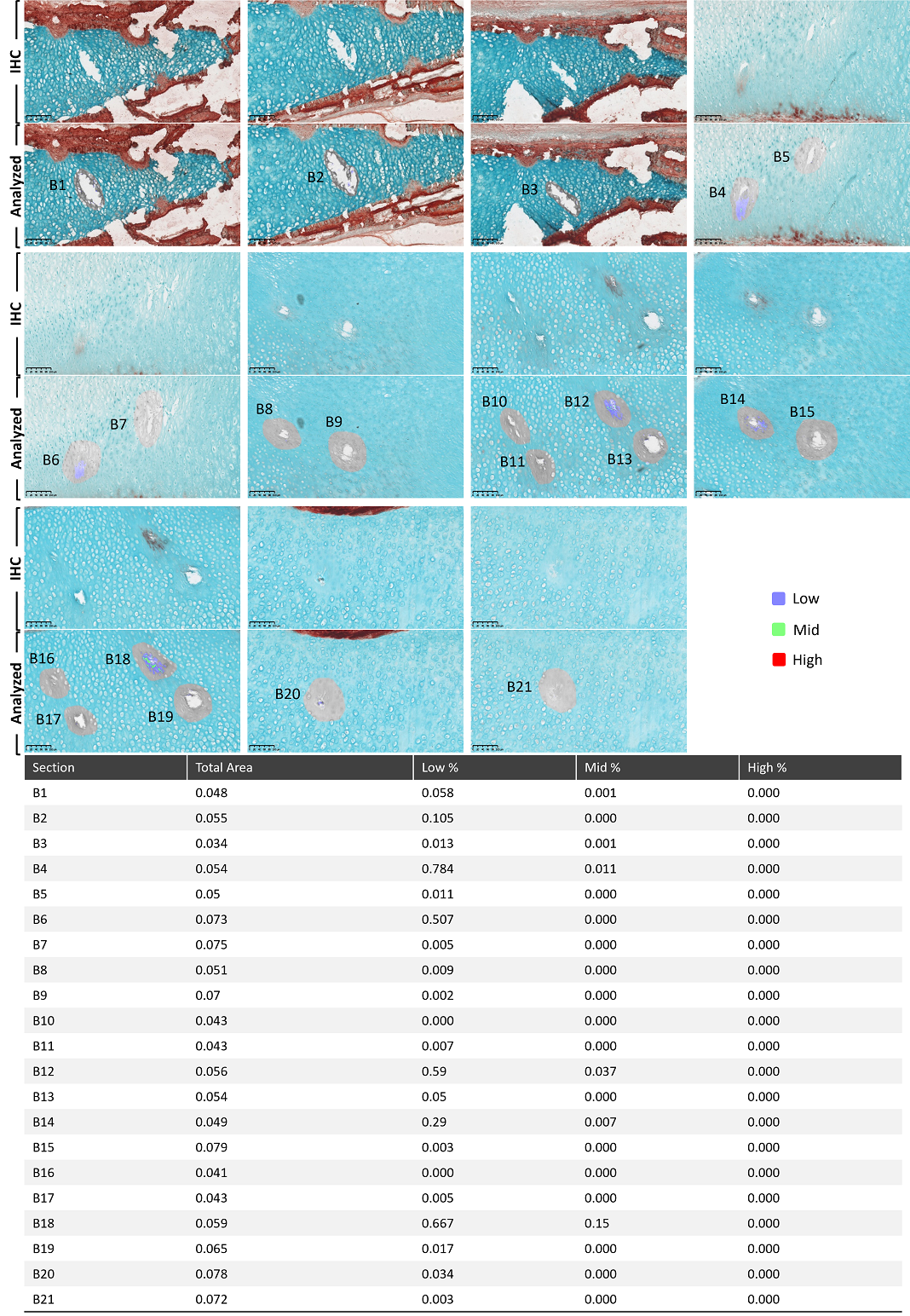

### Supplementary Figure 7

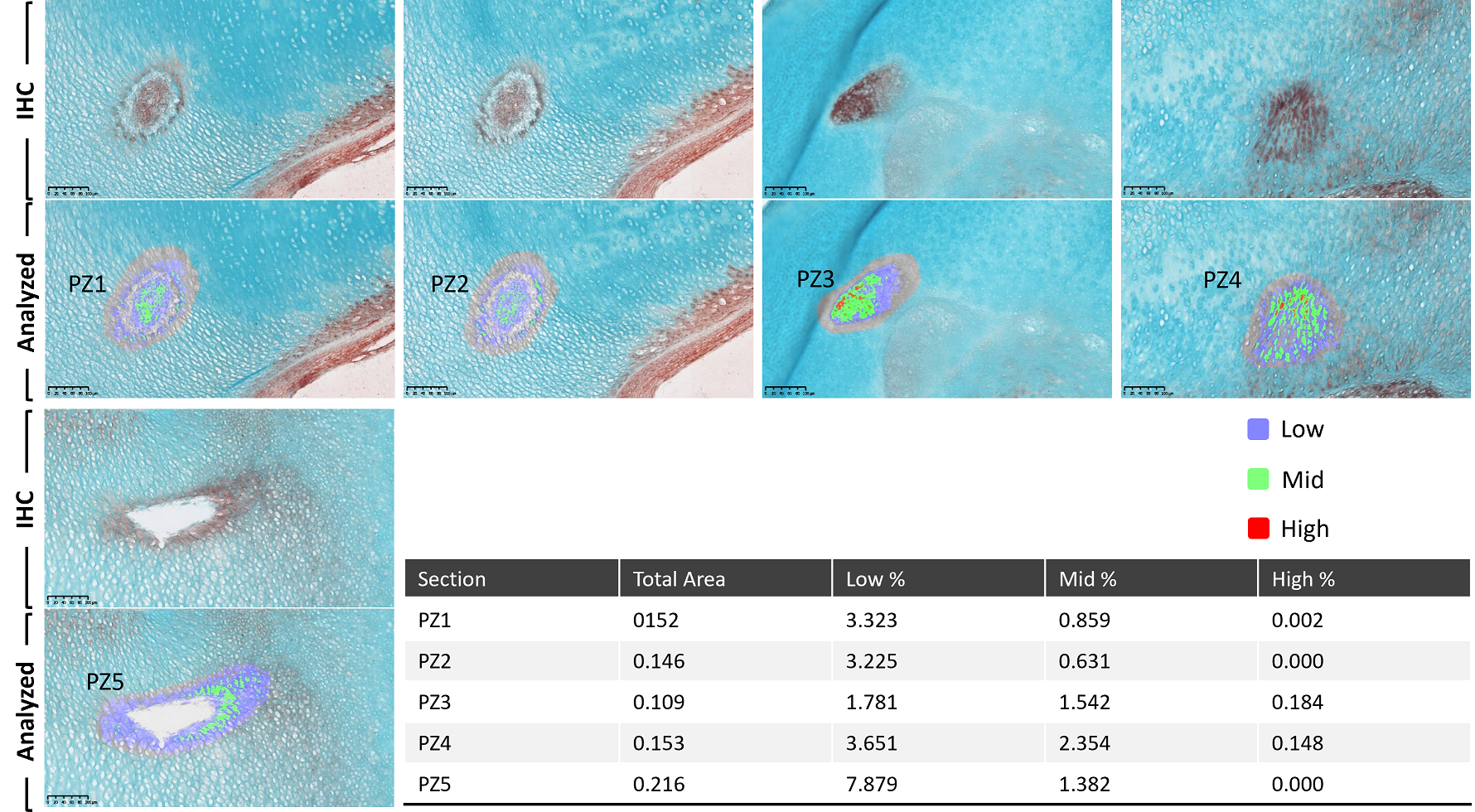

### Supplementary Figure 8

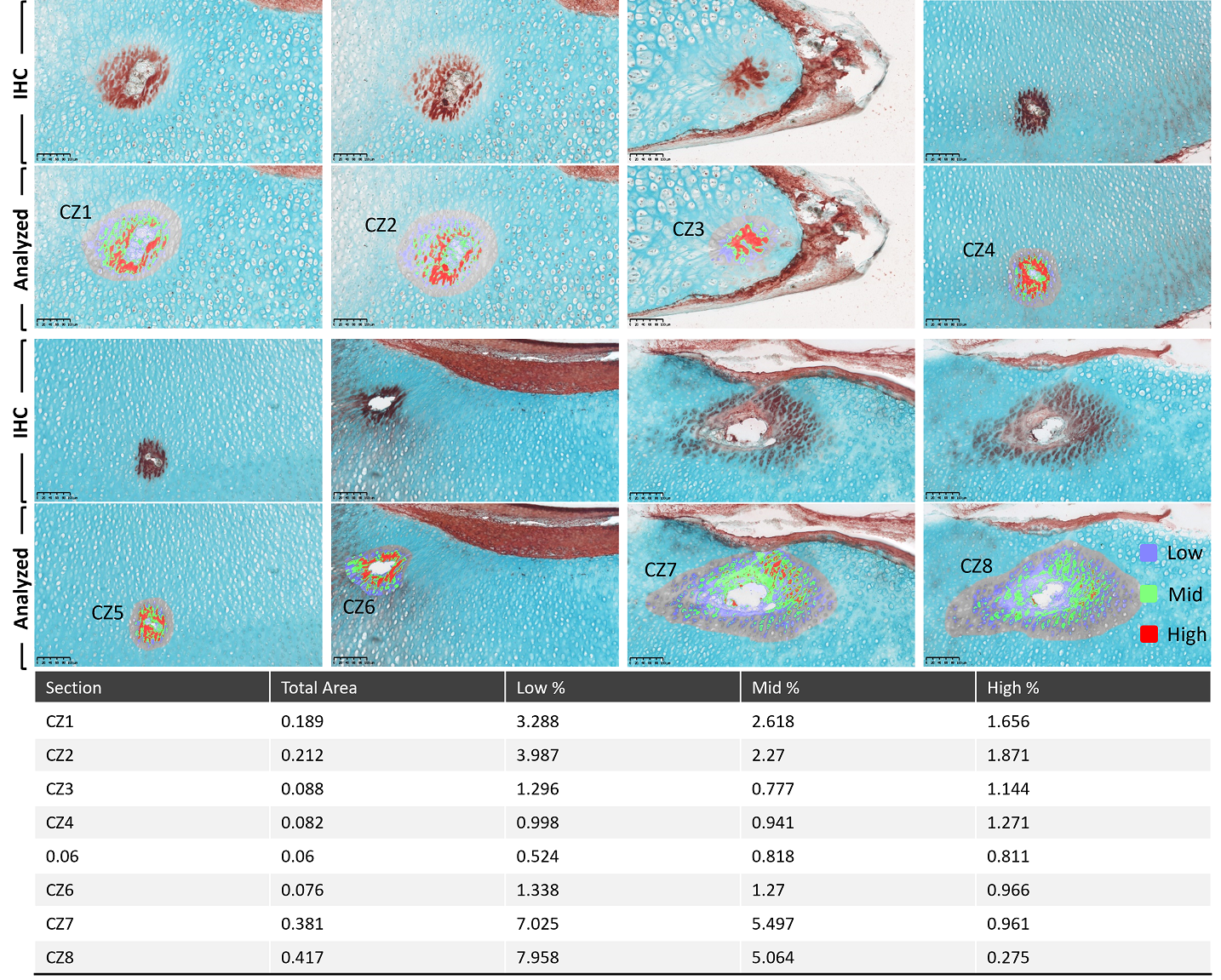
